## Supplementary Note for "Haplotype-based genome wide association study using a novel SNP-set method : RAINBOW"

### Notations

In this section, we define the settings of the following methods, and introduce the frequently used identities.

- $\mathbf{I}_J$ denotes a $J \times J$ identity matrix.

- $\mathbf{1}_{I \times J}$ denotes a $I \times J$ matrix whose elments are all 1.

- MVN $(\boldsymbol{\mu}, \boldsymbol{\Sigma})$ denotes a multivariate normal distribution with a mean $\boldsymbol{\mu}$ and a variance-covariance matrix $\boldsymbol{\Sigma}$.

- For any $a \times b$ matrix $\mathbf{A}$, the $b \times a$ matrix $\mathbf{A}^{\dagger}$ denotes the Moore-Penrose pseudo-inverse of $\mathbf{A}$.

- For any $a \times b$ matrix $\mathbf{A}$ and $a \times b$ matrix $\mathbf{B}$, the $a \times b$ matrix $\mathbf{C} = \mathbf{A} \circ \mathbf{B}$ denotes the Hadamard product between $\mathbf{A}$ and $\mathbf{B}$.

- For any $a \times a$ square matrix $\mathbf{A}$, the expression $|\mathbf{A}|_+$ denotes the pseudo-determinant of $\mathbf{A}$. If $\mathbf{A}$ is positive semi-difinite, $|\mathbf{A}|_+$ will be computed as the product of non-zero eigen values of $\mathbf{A}$.

- $\mathbf{y}$ denotes a $n \times 1$ vector of phenotypic values.

- $\mathbf{X}$ denotes a $n \times p$ covatiate matrix with full column rank $p$ where $n \geq p$.

- $\boldsymbol{\beta}$ denotes a $p \times 1$ vector of fixed effects for covariates.

- $\mathbf{Z}_{\mathrm{c}}$ denotes a $n \times m_{\mathrm{c}}$ design matrix corresponding to random effects $\mathbf{u}_{\mathrm{c}}$ for family relatedness.

- $\mathbf{u}_{\mathrm{c}}$ denotes a $m_{\mathrm{c}} \times 1$ vector of random effects for family relatedness. We assume $\mathbf{u}_{\mathrm{c}} \sim \mathrm{MVN}(\mathbf{0}, \mathbf{K}_{\mathrm{c}}\sigma^2_{\mathrm{c}})$ where $\mathbf{K}_{\mathrm{c}}$ is the additive genetic relationship matrix estimated from marker genotype $\tilde{\mathbf{W}}_{\mathrm{c}}$ and $\sigma^2_{\mathrm{c}}$ is the additive genetic variance.

- $\mathbf{Z}_{\mathrm{r}_i}$ denotes a $n \times m_{\mathrm{r}_i}$ design matrix corresponding to random effects $\mathbf{u}_{\mathrm{r}_i}$ for each SNP-set.

- $\mathbf{u}_{\mathrm{r}_i}$ denotes a $m_{\mathrm{r}_i} \times 1$ vector of random effects for each SNP-set. We assume $\mathbf{u}_{\mathrm{r}_i} \sim \mathrm{MVN}(\mathbf{0}, \mathbf{K}_{\mathrm{r}_i}\sigma^2_{\mathrm{r}_i})$ where $\mathbf{K}_{\mathrm{r}_i}$ is the Gram matrix computed from marker genotype in SNP-set of interest $\tilde{\mathbf{W}}_{\mathrm{r}_i}$ and $\sigma^2_{\mathrm{r}_i}$ is the genetic variance of each SNP-set.

- $\boldsymbol{\epsilon}$ denotes a $n \times 1$ vector of residuals. We assume $\boldsymbol{\epsilon} \sim \mathrm{MVN}(\mathbf{0}, \mathbf{I}_n\sigma^2_{\mathrm{e}})$ where $\sigma^2_{\mathrm{e}}$ is the residual variance.

- $\mathbf{V}$ is a $n \times n$ phenotypic variance-covariance matrix. In this paper, $\mathbf{V} = \mathbf{Z}_{\mathrm{c}}\mathbf{K}_{\mathrm{c}}\mathbf{Z}_{\mathrm{c}}^{\mathrm{T}}\sigma^2_{\mathrm{c}} + \mathbf{Z}_{\mathrm{r}_i}\mathbf{K}_{\mathrm{r}_i}\mathbf{Z}_{\mathrm{r}_i}^{\mathrm{T}}\sigma^2_{\mathrm{r}_i} + \mathbf{I}_n\sigma^2_{\mathrm{e}}$. Under the null hypothesis $\sigma^2_{\mathrm{r}_i} = 0$ since $\sigma^2_{\mathrm{r}_i}$

is the parameter to be tested. $\mathbf{V}$ is assumed to be a full rank $(n)$ matrix.

- $\mathbf{H}$ is a $n \times n$ matrix which satisfies $\mathbf{H} = \mathbf{V}/\sigma_{\mathrm{c}}^2 = \mathbf{Z}_{\mathrm{c}}\mathbf{K}_{\mathrm{c}}\mathbf{Z}_{\mathrm{c}}^{\mathrm{T}} + \mathbf{Z}_{\mathrm{r}_i}\mathbf{K}_{\mathrm{r}_i}\mathbf{Z}_{\mathrm{r}_i}^{\mathrm{T}}\gamma_{\mathrm{r}_i} + \mathbf{I}_n\delta_{\mathrm{e}}$ where $\gamma_{\mathrm{r}_i} = \sigma_{\mathrm{r}_i}^2/\sigma_{\mathrm{c}}^2$ and $\delta_{\mathrm{e}} = \sigma_{\mathrm{e}}^2/\sigma_{\mathrm{c}}^2$ to be estimated.

- $\mathbf{S} = \mathbf{I}_n - \mathbf{X}\left(\mathbf{X}^{\mathrm{T}}\mathbf{X}\right)^{-1}\mathbf{X}^{\mathrm{T}}$ is a $n \times n$ symmetric covariance orthogonal projection matrix with rank $n - p$.

- $\mathbf{P} = \mathbf{I}_n - \mathbf{X}\left(\mathbf{X}^{\mathrm{T}}\mathbf{V}^{-1}\mathbf{X}\right)^{-1}\mathbf{X}^{\mathrm{T}}\mathbf{V}^{-1} = \mathbf{I}_n - \mathbf{X}\left(\mathbf{X}^{\mathrm{T}}\mathbf{H}^{-1}\mathbf{X}\right)^{-1}\mathbf{X}^{\mathrm{T}}\mathbf{H}^{-1}$ is a $n \times n$ matrix with rank $n - p$.

- $\mathbf{w} = \mathbf{P}\mathbf{y}$ is a $n \times 1$ vector used for the REML estimation. We also define $\mathbf{w}_{\mathrm{L}}$, $\mathbf{w}_{\mathrm{R}}$, $\mathbf{P}_{\mathrm{L}}$ and $\mathbf{P}_{\mathrm{R}}$ by

$$\mathbf{w} = \begin{bmatrix} \mathbf{w}_{\mathrm{L}} \\ \mathbf{w}_{\mathrm{R}} \end{bmatrix} = \begin{bmatrix} \mathbf{P}_{\mathrm{L}} \\ \mathbf{P}_{\mathrm{R}} \end{bmatrix} \mathbf{y} = \mathbf{P}\mathbf{y},$$

  where $\mathbf{w}_{\mathrm{L}}$ (or $\mathbf{w}_{\mathrm{R}}$) stands for a linearly independent (or redundant) part of $\mathbf{w}$, and $\mathbf{P}_{\mathrm{L}}$ (or $\mathbf{P}_{\mathrm{R}}$) denotes the corresponding partition of $\mathbf{P}$. Here, $\mathbf{P}_{\mathrm{L}}$ should satisfy $\mathbf{P}_{\mathrm{L}}^{\mathrm{T}}\mathbf{P}_{\mathrm{L}} = \mathbf{S}$ and $\mathbf{P}_{\mathrm{L}}\mathbf{P}_{\mathrm{L}}^{\mathrm{T}} = \mathbf{I}_{n-p}$ [1, 2, 3]. We also define the eigen decomposition of $\mathbf{SHS}$ as

$$\mathbf{SHS} = \mathbf{U}_{\mathrm{L}}\mathbf{\Lambda}_L\mathbf{U}_{\mathrm{L}}^{\mathrm{T}},$$

  where $\mathbf{\Lambda}_L$ is a $n-p \times n-p$ diagonal matrix whose elements are non-zero eigen values of $\mathbf{SHS}$ in the decreasing order, and $\mathbf{U}_{\mathrm{L}}$ is a $n \times n - p$ eigen vector matrix whose each eigen vector corresponds to each eigen value. Here, $\mathbf{U}_{\mathrm{L}}$ is the part of the unitary matrix with first $n - p$ columns of eigen vectors, and it satisfies $\mathbf{U}_{\mathrm{L}}\mathbf{U}_{\mathrm{L}}^{\mathrm{T}} = \mathbf{S}$ and $\mathbf{U}_{\mathrm{L}}^{\mathrm{T}}\mathbf{U}_{\mathrm{L}} = \mathbf{I}_{n-p}$ (Proposition 7). Therefore, we can say $\mathbf{P}_{\mathrm{L}} = \mathbf{U}_{\mathrm{L}}^{\mathrm{T}}$.

- $\mathbf{Q} = \mathbf{H}^{-1} - \mathbf{H}^{-1}\mathbf{X}\left(\mathbf{X}^{\mathrm{T}}\mathbf{H}^{-1}\mathbf{X}\right)^{-1}\mathbf{X}^{\mathrm{T}} = \mathbf{H}^{-1}\mathbf{P} = \mathbf{P}^{\mathrm{T}}\mathbf{H}^{-1}$ is a $n \times n$ symmetry matrix with rank $n - p$.

- $\mathbf{W}_{\mathrm{r}_i}$ is a $m_{\mathrm{r}_i} \times M_i$ mmarker genotype matrix belonging to the $i_{\mathrm{th}}$ SNP-set. Here, $M_i$ is the number of SNPs in the $i_{\mathrm{th}}$ SNP-set.

### Multi-kernel linear mixed model

In this study, the alternative model can be written as the multi-kernel mixed model.

$$\mathbf{y} = \mathbf{X}\boldsymbol{\beta} + \mathbf{Z}_{\mathrm{c}}\mathbf{u}_{\mathrm{c}} + \mathbf{Z}_{\mathrm{r}_i}\mathbf{u}_{\mathrm{r}_i} + \boldsymbol{\epsilon} \tag{1}$$

On the other hand, the null model can be written as

$$\mathbf{y} = \mathbf{X}\boldsymbol{\beta} + \mathbf{Z}_{\mathrm{c}}\mathbf{u}_{\mathrm{c}} + \boldsymbol{\epsilon} \tag{2}$$

Therefore, we somehow test the null hypothesis $H_0 : \sigma_{\mathrm{r}_i}^2 = 0$ for evaluating the significance of the effects of SNP-set of interst.

### Restricted maximum likelihood (REML)

For the multi-kernel linear mixed model described above, the restricted log likelihood of $\mathbf{y}$ can be regarded as the log likelihood of $\mathbf{w}_{\mathrm{L}}$, and it can be expressed as a quite simple format by only using $\mathbf{Q}$.

$$\begin{aligned} l_{\mathrm{R}}(\mathbf{y};\sigma_{\mathrm{c}},\gamma_{\mathrm{r}_i},\delta_{\mathrm{e}}) &= -\frac{n-p}{2}\log(2\pi) - \frac{1}{2}\log\left|\mathbf{P}_{\mathrm{L}}\mathbf{V}\mathbf{P}_{\mathrm{L}}^{\mathrm{T}}\right| - \frac{1}{2}\mathbf{y}\mathbf{P}_{\mathrm{L}}^{\mathrm{T}}\left(\mathbf{P}_{\mathrm{L}}\mathbf{V}\mathbf{P}_{\mathrm{L}}^{\mathrm{T}}\right)^{-1}\mathbf{P}_{\mathrm{L}}\mathbf{y} \\ &= -\frac{n-p}{2}\log(2\pi) + \frac{1}{2}\log\left|\mathbf{P}^{\mathrm{T}}\mathbf{V}^{-1}\mathbf{P}\right|_{+} - \frac{1}{2}\mathbf{y}^{\mathrm{T}}\mathbf{V}^{-1}\mathbf{P}\mathbf{y} \\ &= -\frac{n-p}{2}\log(2\pi\sigma_{\mathrm{c}}^2) + \frac{1}{2}\log|\mathbf{Q}|_{+} - \frac{1}{2\sigma_{\mathrm{c}}^2}\mathbf{y}^{\mathrm{T}}\mathbf{Q}\mathbf{y} \qquad (3) \end{aligned}$$

by using Proposition 8 and 9.
In addition, by using Proposition 10, this restricted log likelihood is same as the well known format [4, 5, 6] as follows.

$$\begin{aligned} l_{\mathrm{R}}(\mathbf{y};\sigma_{\mathrm{c}},\gamma_{\mathrm{r}_i},\delta_{\mathrm{e}}) &= -\frac{n-p}{2}\log(2\pi\sigma_{\mathrm{c}}^2) + \frac{1}{2}\log|\mathbf{Q}|_{+} - \frac{1}{2\sigma_{\mathrm{c}}^2}\mathbf{y}^{\mathrm{T}}\mathbf{Q}\mathbf{y} \\ &= -\frac{n-p}{2}\log(2\pi\sigma_{\mathrm{c}}^2) - \frac{1}{2}\log|\mathbf{H}| + \frac{1}{2}\log\left|\mathbf{X}^{\mathrm{T}}\mathbf{X}\right| \\ &\quad - \frac{1}{2}\log\left|\mathbf{X}^{\mathrm{T}}\mathbf{H}^{-1}\mathbf{X}\right| - \frac{1}{2\sigma_{\mathrm{c}}^2}\mathbf{y}^{\mathrm{T}}\mathbf{Q}\mathbf{y} \qquad (4) \end{aligned}$$

Plugging $\hat{\sigma}_{\mathrm{c}}^2 = \mathbf{y}^{\mathrm{T}}\mathbf{Q}\mathbf{y}/(n-p)$ imto Eq. (3), we get

$$l_{\mathrm{R}}(\mathbf{y};\hat{\sigma}_{\mathrm{c}},\gamma_{\mathrm{r}_i},\delta_{\mathrm{e}}) = -\frac{n-p}{2}\left\{\log\left(\frac{2\pi e}{n-p}\right) + \log\left(\mathbf{y}^{\mathrm{T}}\mathbf{Q}\mathbf{y}\right)\right\} + \frac{1}{2}\log|\mathbf{Q}|_{+} \qquad (5)$$

### Efficient likelihood ratio test used in RAINBOW

In this section, we describe how to implenment computationally efficient algorithm for the likelihood ratio (LR) test for the two kernel linear mixed model.

In this study, we assume the Gram matrix for each SNP-set $\mathbf{Z}_{\mathrm{r}_i}\mathbf{K}_{\mathrm{r}_i}\mathbf{Z}_{\mathrm{r}_i}^{\mathrm{T}}$ is low rank, so $rank(\mathbf{Z}_{\mathrm{r}_i}\mathbf{K}_{\mathrm{r}_i}\mathbf{Z}_{\mathrm{r}_i}^{\mathrm{T}}) \ll n$. Then, $\mathbf{H}$ can be written as

$$\begin{aligned} \mathbf{H} &= \mathbf{Z}_{\mathrm{c}}\mathbf{K}_{\mathrm{c}}\mathbf{Z}_{\mathrm{c}}^{\mathrm{T}} + \mathbf{Z}_{\mathrm{r}_i}\mathbf{K}_{\mathrm{r}_i}\mathbf{Z}_{\mathrm{r}_i}^{\mathrm{T}}\gamma_{\mathrm{r}_i} + \mathbf{I}_n\delta_{\mathrm{e}} \\ &= \mathbf{Z}_{\mathrm{c}}\mathbf{K}_{\mathrm{c}}\mathbf{Z}_{\mathrm{c}}^{\mathrm{T}} + \tilde{\mathbf{W}}_{\mathrm{r}_i}\tilde{\mathbf{\Gamma}}_{\mathrm{r}_i}\tilde{\mathbf{W}}_{\mathrm{r}_i}^{\mathrm{T}} + \mathbf{I}_n\delta_{\mathrm{e}}, \qquad (6) \end{aligned}$$

where $\tilde{\mathbf{W}}_{\mathrm{r}_i}$ is a $n \times k$ matrix and $\tilde{\mathbf{\Gamma}}_{\mathrm{r}_i}$ is a $k \times k$ square matrix. Here, $k$ is the rank of $\mathbf{Z}_{\mathrm{r}_i}\mathbf{K}_{\mathrm{r}_i}\mathbf{Z}_{\mathrm{r}_i}^{\mathrm{T}}$, so $k \ll n$. Concrete examples of $\tilde{\mathbf{W}}_{\mathrm{r}_i}$ and $\tilde{\mathbf{\Gamma}}_{\mathrm{r}_i}$ will be described later.

#### Low rank update of $\mathbf{Q}$

One of the drawbacks of the LR test is a large amount of computation because the LR test requires the maximization of restricted likelihood for each SNP-set. To reduce the computational complexity, [7, 8] proposed the efficient computation of the restricted log likelihood Eq. (5) by using low rank update of $\mathbf{Q}$ as follows.

First, the low-rank update of $\mathbf{Q}$ is

$$
\begin{aligned}
\mathbf{Q} &= \mathbf{P}^{\mathrm{T}}\mathbf{H}^{-1}\mathbf{P} \\
&= (\mathbf{SHS})^{\dagger} \\
&= \left(\mathbf{S}\left(\mathbf{Z}_{\mathrm{c}}\mathbf{K}_{\mathrm{c}}\mathbf{Z}_{\mathrm{c}}^{\mathrm{T}} + \tilde{\mathbf{W}}_{\mathrm{r}_i}\tilde{\mathbf{\Gamma}}_{\mathrm{r}_i}\tilde{\mathbf{W}}_{\mathrm{r}_i}^{\mathrm{T}} + \mathbf{I}_n\delta_{\mathrm{e}}\right)\mathbf{S}\right)^{\dagger} \\
&= \left(\mathbf{S}\left(\mathbf{Z}_{\mathrm{c}}\mathbf{K}_{\mathrm{c}}\mathbf{Z}_{\mathrm{c}}^{\mathrm{T}} + \mathbf{I}_n\delta_{\mathrm{e}}\right)\mathbf{S} + \mathbf{S}\tilde{\mathbf{W}}_{\mathrm{r}_i}\tilde{\mathbf{\Gamma}}_{\mathrm{r}_i}\tilde{\mathbf{W}}_{\mathrm{r}_i}^{\mathrm{T}}\mathbf{S}\right)^{\dagger} \\
&= \left(\mathbf{U}_{\mathrm{L}}\left(\mathbf{\Lambda}_{\mathrm{c}} + \mathbf{I}_{n-p}\delta_{\mathrm{e}}\right)\mathbf{U}_{\mathrm{L}}^{\mathrm{T}} + \mathbf{U}_{\mathrm{L}}\mathbf{U}_{\mathrm{L}}^{\mathrm{T}}\tilde{\mathbf{W}}_{\mathrm{r}_i}\tilde{\mathbf{\Gamma}}_{\mathrm{r}_i}\tilde{\mathbf{W}}_{\mathrm{r}_i}^{\mathrm{T}}\mathbf{U}_{\mathrm{L}}\mathbf{U}_{\mathrm{L}}^{\mathrm{T}}\right)^{\dagger} \\
&= \left(\mathbf{U}_{\mathrm{L}}\left(\mathbf{\Lambda}_{\mathrm{c}} + \mathbf{I}_{n-p}\delta_{\mathrm{e}} + \mathbf{U}_{\mathrm{L}}^{\mathrm{T}}\tilde{\mathbf{W}}_{\mathrm{r}_i}\tilde{\mathbf{\Gamma}}_{\mathrm{r}_i}\tilde{\mathbf{W}}_{\mathrm{r}_i}^{\mathrm{T}}\mathbf{U}_{\mathrm{L}}\right)\mathbf{U}_{\mathrm{L}}^{\mathrm{T}}\right)^{\dagger} \\
&= \mathbf{U}_{\mathrm{L}}\left(\mathbf{\Lambda}_{\mathrm{c}} + \mathbf{I}_{n-p}\delta_{\mathrm{e}} + \mathbf{U}_{\mathrm{L}}^{\mathrm{T}}\tilde{\mathbf{W}}_{\mathrm{r}_i}\tilde{\mathbf{\Gamma}}_{\mathrm{r}_i}\tilde{\mathbf{W}}_{\mathrm{r}_i}^{\mathrm{T}}\mathbf{U}_{\mathrm{L}}\right)^{-1}\mathbf{U}_{\mathrm{L}}^{\mathrm{T}} \\
&= \mathbf{U}_{\mathrm{L}}\left(\mathbf{\Lambda}_{\mathrm{c}} + \mathbf{I}_{n-p}\delta_{\mathrm{e}}\right)^{-1}\mathbf{U}_{\mathrm{L}}^{\mathrm{T}} \\
&\quad - \mathbf{U}_{\mathrm{L}}\left(\mathbf{\Lambda}_{\mathrm{c}} + \mathbf{I}_{n-p}\delta_{\mathrm{e}}\right)^{-1}\mathbf{U}_{\mathrm{L}}^{\mathrm{T}}\tilde{\mathbf{W}}_{\mathrm{r}_i} \\
&\quad \cdot \left(\tilde{\mathbf{\Gamma}}_{\mathrm{r}_i}^{-1} + \tilde{\mathbf{W}}_{\mathrm{r}_i}^{\mathrm{T}}\mathbf{U}_{\mathrm{L}}\left(\mathbf{\Lambda}_{\mathrm{c}} + \mathbf{I}_{n-p}\delta_{\mathrm{e}}\right)^{-1}\mathbf{U}_{\mathrm{L}}^{\mathrm{T}}\tilde{\mathbf{W}}_{\mathrm{r}_i}\right)^{-1} \\
&\quad \cdot \tilde{\mathbf{W}}_{\mathrm{r}_i}^{\mathrm{T}}\mathbf{U}_{\mathrm{L}}\left(\mathbf{\Lambda}_{\mathrm{c}} + \mathbf{I}_{n-p}\delta_{\mathrm{e}}\right)^{-1}\mathbf{U}_{\mathrm{L}}^{\mathrm{T}} \\
&= \mathbf{O}_{\mathrm{c}} - \mathbf{O}_{\mathrm{c}}\tilde{\mathbf{W}}_{\mathrm{r}_i}\left(\tilde{\mathbf{\Gamma}}_{\mathrm{r}_i}^{-1} + \tilde{\mathbf{W}}_{\mathrm{r}_i}^{\mathrm{T}}\mathbf{O}_{\mathrm{c}}\tilde{\mathbf{W}}_{\mathrm{r}_i}\right)^{-1}\tilde{\mathbf{W}}_{\mathrm{r}_i}^{\mathrm{T}}\mathbf{O}_{\mathrm{c}}
\end{aligned} \tag{7}
$$

Here, we use Proposition 6, 5, 2, 7 and the Woodbury identity, and we define the eigen decomposition of $\mathbf{S}\mathbf{Z}_{\mathrm{c}}\mathbf{K}_{\mathrm{c}}\mathbf{Z}_{\mathrm{c}}^{\mathrm{T}}\mathbf{S} = \mathbf{U}_{\mathrm{L}}\mathbf{\Lambda}_{\mathrm{c}}\mathbf{U}_{\mathrm{L}}^{\mathrm{T}}$ where $\mathbf{\Lambda}_{\mathrm{c}}$ is a $n-p \times n-p$ diagonal matrix and $n \times n$ matrix $\mathbf{O}_{\mathrm{c}} = \mathbf{U}_{\mathrm{L}}\left(\mathbf{\Lambda}_{\mathrm{c}} + \mathbf{I}_{n-p}\delta_{\mathrm{e}}\right)^{-1}\mathbf{U}_{\mathrm{L}}^{\mathrm{T}}$ to shorten the notation. Assuming that the eigen decomposition of $\mathbf{S}\left(\mathbf{Z}_{\mathrm{c}}\mathbf{K}_{\mathrm{c}}\mathbf{Z}_{\mathrm{c}}^{\mathrm{T}} + \mathbf{I}_{n-p}\delta_{\mathrm{e}}\right)\mathbf{S}$ has been pre-computed, the additional computation required will be $\mathbf{O}_{\mathrm{c}}\tilde{\mathbf{W}}_{\mathrm{r}_i}$, an $O(n^2k)$ operation.

*Update of the squared form* $\mathbf{y}^{\mathrm{T}}\mathbf{Q}\mathbf{y}$

Since we know the derivation of the low-rank update of $\mathbf{Q}$ (Eq. (7)) now, we can plug this into the squared form.

$$
\mathbf{y}^{\mathrm{T}}\mathbf{Q}\mathbf{y} = \mathbf{y}^{\mathrm{T}}\mathbf{O}_{\mathrm{c}}\mathbf{y} - \mathbf{y}^{\mathrm{T}}\mathbf{O}_{\mathrm{c}}\tilde{\mathbf{W}}_{\mathrm{r}_i}\left(\tilde{\mathbf{\Gamma}}_{\mathrm{r}_i}^{-1} + \tilde{\mathbf{W}}_{\mathrm{r}_i}^{\mathrm{T}}\mathbf{O}_{\mathrm{c}}\tilde{\mathbf{W}}_{\mathrm{r}_i}\right)^{-1}\tilde{\mathbf{W}}_{\mathrm{r}_i}^{\mathrm{T}}\mathbf{O}_{\mathrm{c}}\mathbf{y} \tag{8}
$$

*Update of the determinant* $\log|\mathbf{Q}|_+$

Since the update matrix is not necessarily positive semi-definite, we have to slightly modify Eq. (7) to avoid numerical instability.

$$
\begin{aligned}
\log|\mathbf{Q}|_+ &= -\log\left|\mathbf{U}_{\mathrm{L}}^{\mathrm{T}}\mathbf{H}\mathbf{U}_{\mathrm{L}}\right| \\
&= -\log\left|\mathbf{I}_{n-p}\delta_{\mathrm{e}} + \mathbf{\Lambda}_{\mathrm{c}} + \mathbf{U}_{\mathrm{L}}^{\mathrm{T}}\tilde{\mathbf{W}}_{\mathrm{r}_i}\tilde{\mathbf{\Gamma}}_{\mathrm{r}_i}\tilde{\mathbf{W}}_{\mathrm{r}_i}^{\mathrm{T}}\mathbf{U}_{\mathrm{L}}\right| \\
&= -\log\left|\mathbf{I}_{n-p}\delta_{\mathrm{e}} + \mathbf{\Lambda}_{\mathrm{c}} + \mathbf{AB}\right| \\
&= -\log\left(\left|\mathbf{I}_{n-p}\delta_{\mathrm{e}} + \mathbf{\Lambda}_{\mathrm{c}}\right| \cdot \left|\mathbf{I}_{n-p} + \left(\mathbf{I}_{n-p}\delta_{\mathrm{e}} + \mathbf{\Lambda}_{\mathrm{c}}\right)^{-1}\mathbf{AB}\right|\right) \\
&= -\log\left|\mathbf{I}_{n-p}\delta_{\mathrm{e}} + \mathbf{\Lambda}_{\mathrm{c}}\right| - \log\left|\mathbf{I}_k + \mathbf{B}\left(\mathbf{I}_{n-p}\delta_{\mathrm{e}} + \mathbf{\Lambda}_{\mathrm{c}}\right)^{-1}\mathbf{A}\right|
\end{aligned} \tag{9}
$$

Here, we used Proposition 5, 6, 7 and the Sylvester's determinant identity, and we also define a $n-p \times k$ matrix $\mathbf{A} = \mathbf{U}_{\mathrm{L}}^{\mathrm{T}} \tilde{\mathbf{W}}_{\mathrm{r}_i} \tilde{\mathbf{\Gamma}}_{\mathrm{r}_i}$ and a $k \times n-p$ matrix $\mathbf{B} = \tilde{\mathbf{W}}_{\mathrm{r}_i}^{\mathrm{T}} \mathbf{U}_{\mathrm{L}}$. When calculating the second term of Eq. (9), we have to be careful about the possibility that $\mathbf{I}_k + \mathbf{B} \left( \mathbf{I}_{n-p} \delta_{\mathrm{e}} + \mathbf{\Lambda}_{\mathrm{c}} \right)^{-1} \mathbf{A}$ is not necessarily positive semi-definite. Since the matrix $\mathbf{Q}$ is positive semi-definite, it is possible to calculate the second term of Eq. (9), however, the matrix $\mathbf{I}_k + \mathbf{B} \left( \mathbf{I}_{n-p} \delta_{\mathrm{e}} + \mathbf{\Lambda}_{\mathrm{c}} \right)^{-1} \mathbf{A}$ has an even numbers of negative eigen values, so instead of simply computing the determinant of this matrix, we have to avoid taking logarithms of such negative eigen values.

#### Estimation of variance components

Now we can derive the efficient computation of the restricted log likelihood from Eq. (5), Eq. (8) and Eq. (9), we can optimize $\gamma_{\mathrm{r}_i}$ and $\delta_{\mathrm{e}}$ over maximazation of Eq. (5) by using L-BFGS optimzation method as we introduce in the paper [9].

Then, we calculated the weighted variance-covariance matrix by $\gamma_{\mathrm{r}_i}$, and reestimate variance components by using EMMA (efficient mixed model association) or GEMMA (genome wide efficient mixed model association) [6, 10].

#### Discussion on the identity of $\tilde{\mathbf{W}}_{\mathrm{r}_i}$ and $\tilde{\mathbf{\Gamma}}_{\mathrm{r}_i}$

In this subsection, we discuss on what $\tilde{\mathbf{W}}_{\mathrm{r}_i}$ and $\tilde{\mathbf{\Gamma}}_{\mathrm{r}_i}$ correspond to depending on the kind of the kernels for $\mathbf{K}_{\mathrm{r}_i}$.

*For the case where $\mathbf{K}_{\mathrm{r}_i}$ is the linear kernel*

Here, we discuss the case where $\mathbf{K}_{\mathrm{r}_i}$ is calculated as the linear kernel of the marker genotype $\mathbf{W}_{\mathrm{r}_i}$ belonging to the $i$ th SNP-set.

For example, if we assume $\mathbf{K}_{\mathrm{r}_i}$ is the additive genetic matrix of $\mathbf{W}_{\mathrm{r}_i}$, $\tilde{\mathbf{W}}_{\mathrm{r}_i}$ corresponds to

$$\tilde{\mathbf{W}}_{\mathrm{r}_i} = \frac{\mathbf{Z}_{\mathrm{r}_i} \left( \mathbf{W}_{\mathrm{r}_i} + \mathbf{1}_{m_{\mathrm{r}_i} \times M_i} - 2 \cdot \mathbf{\Phi} \right)}{2 \cdot \sum_{m=1}^{M_i} p_m \left( 1 - p_m \right)}, \tag{10}$$

where $p_m$ is the allele frequency of the 1 allele at marker $m$ and $\mathbf{\Phi}$ is a $m_{\mathrm{r}_i} \times M_i$ matrix whose $m_{\mathrm{th}}$ column equals to $\mathbf{\Phi} = p_m \cdot \mathbf{1}_{m_{\mathrm{r}_i} \times 1}$ [11].

On the other hand, $\tilde{\mathbf{\Gamma}}_{\mathrm{r}_i}$ corresponds to

$$\tilde{\mathbf{\Gamma}}_{\mathrm{r}_i} = \mathbf{I}_{M_i} \gamma_{\mathrm{r}_i}, \tag{11}$$

so in this case, $\tilde{\mathbf{\Gamma}}_{\mathrm{r}_i}$ is a $M_i \times M_i$ diagonal matrix. Here, $k$ in Eq. (7) equals to the number of SNPs in the $i$ th SNP-set, $M_i$.

*For the case where $\mathbf{K}_{\mathrm{r}_i}$ is the exponential or gaussian kernel*

Here, we discuss the case where $\mathbf{K}_{\mathrm{r}_i}$ is calculated as the exponential or gaussian kernel of the marker genotype $\mathbf{W}_{\mathrm{r}_i}$ belonging to the $i$ th SNP-set.

First, we calculated the Euclidean distance matrix $\mathbf{D}_{\mathrm{r}_i}$ from the marker genotype $\mathbf{W}_{\mathrm{r}_i}$ belonging to the $i$ th SNP-set. Then for the case where $\mathbf{K}_{\mathrm{r}_i}$ is the exponential kernel, $\mathbf{K}_{\mathrm{r}_i}$ is calculated as

$$\mathbf{K}_{\mathrm{r}_i} = \exp\left( -\frac{h_{\mathrm{r}_i} \mathbf{D}_{\mathrm{r}_i}}{\sqrt{M_i}} \right), \tag{12}$$

where $h_{\mathrm{r}_i}$ is a hyperparameter calculated as the inverse of median of the off-diagonal elements of $\mathbf{D}^2_{\mathrm{r}_i}/M_i$ for the default setting of RAINBOW. To scale the distance matrix, the distance matrix is divided by $\sqrt{M_i}$ in the exponential. Similarly, for the case where $\mathbf{K}_{\mathrm{r}_i}$ is the gaussian kernel, $\mathbf{K}_{\mathrm{r}_i}$ is calculated as

$$\mathbf{K}_{\mathrm{r}_i} = \exp\left(-\frac{h_{\mathrm{r}_i}\mathbf{D}^2_{\mathrm{r}_i}}{M_i}\right), \tag{13}$$

where the term in the exponential is also deivide by $M_i$ to fit its scale to the linear kernel.

In this case, we cannot apply the decomposition of $\mathbf{K}_{\mathrm{r}_i}$ as seen in the linear kernel case, however, it is assumed that the rank of $\mathbf{K}_{\mathrm{r}_i}$ is still much smaller than the number of observations, $n$. This is because, since there should be strong linkage disequilibrium between SNPs in each SNP-set and many accessions share the same SNPs in that SNP-set, the number of genotypes in that SNP-set $m_{\mathrm{r}_i}$ may be much smaller than $n$. Therefore, in such case,

$$\tilde{\mathbf{W}}_{\mathrm{r}_i} = \mathbf{Z}_{\mathrm{r}_i}, \tag{14}$$

$$\tilde{\mathbf{\Gamma}}_{\mathrm{r}_i} = \mathbf{K}_{\mathrm{r}_i}\gamma_{\mathrm{r}_i} \tag{15}$$

Here, $k$ in Eq. (7) equals to the number of SNPs in the genotypes in the $i_{\mathrm{th}}$ SNP-set, $m_{\mathrm{r}_i}$. We should be careful about that $\tilde{\mathbf{\Gamma}}_{\mathrm{r}_i}$ is not a diagonal matrix in this case.

### Efficient likelihood ratio test for dominance and epistatic effects

In this section, we extend the efficient LR test for the one random effect of each SNP-set to that for multiple random effects of the SNP-set. In this case, the multi-kernel mixed model will be

$$\mathbf{y} = \mathbf{X}\boldsymbol{\beta} + \mathbf{Z}_{\mathrm{c}}\mathbf{u}_{\mathrm{c}} + \sum_{l=1}^{L}\mathbf{Z}_{\mathrm{r}_i,l}\mathbf{u}_{\mathrm{r}_i,l} + \boldsymbol{\epsilon}, \tag{16}$$

where $\mathbf{u}_{\mathrm{r}_i,l}$ is the $l_{\mathrm{th}}$ random effect of the $i_{\mathrm{th}}$ SNP-set and $\mathbf{Z}_{\mathrm{r}_i,l}$ is a $n \times m_{\mathrm{r}_i,l}$ design matrix which correspond to $\mathbf{u}_{\mathrm{r}_i,l}$. Examples of multiple random effects are additive effects, dominance effects and epistatic effects, and the test for the significance of these effects will be described later.

The restricted log likelihood is still Eq. (5), however, here

$$\begin{aligned}\mathbf{H} &= \mathbf{Z}_{\mathrm{c}}\mathbf{K}_{\mathrm{c}}\mathbf{Z}_{\mathrm{c}}^{\mathrm{T}} + \sum_{l=1}^{L}\mathbf{Z}_{\mathrm{r}_i,l}\mathbf{K}_{\mathrm{r}_i,l}\mathbf{Z}_{\mathrm{r}_i,l}^{\mathrm{T}}\gamma_{\mathrm{r}_i,l} + \mathbf{I}_n\delta_{\mathrm{e}} \\ &= \mathbf{Z}_{\mathrm{c}}\mathbf{K}_{\mathrm{c}}\mathbf{Z}_{\mathrm{c}}^{\mathrm{T}} + \sum_{l=1}^{L}\tilde{\mathbf{W}}_{\mathrm{r}_i,l}\tilde{\mathbf{\Gamma}}_{\mathrm{r}_i,l}\tilde{\mathbf{W}}_{\mathrm{r}_i,l}^{\mathrm{T}} + \mathbf{I}_n\delta_{\mathrm{e}},\end{aligned} \tag{17}$$

where $\mathbf{K}_{\mathrm{r}_i,l}$, $\gamma_{\mathrm{r}_i,l}$, $\tilde{\mathbf{W}}_{\mathrm{r}_i,l}$ and $\tilde{\mathbf{\Gamma}}_{\mathrm{r}_i,l}$ are the extensions of $\mathbf{K}_{\mathrm{r}_i}$, $\gamma_{\mathrm{r}_i}$, $\tilde{\mathbf{W}}_{\mathrm{r}_i}$ and $\tilde{\mathbf{\Gamma}}_{\mathrm{r}_i}$ for the $l_{\mathrm{th}}$ random effect.

#### Low rank update of $\mathbf{Q}^{(L)}$

In this case, the low rank update of $\mathbf{Q}$ is realized by extending Eq. (7) to

$$
\begin{aligned}
\mathbf{Q}^{(L)} &= \mathbf{P}^{\mathrm{T}}\mathbf{H}^{-1}\mathbf{P} \\
&= (\mathbf{SHS})^{\dagger} \\
&= \left(\mathbf{S}\left(\mathbf{Z}_{\mathrm{c}}\mathbf{K}_{\mathrm{c}}\mathbf{Z}_{\mathrm{c}}^{\mathrm{T}} + \sum_{l=1}^{L}\tilde{\mathbf{W}}_{\mathrm{r}_i,l}\tilde{\mathbf{\Gamma}}_{\mathrm{r}_i,l}\tilde{\mathbf{W}}_{\mathrm{r}_i,l}^{\mathrm{T}} + \mathbf{I}_n\delta_{\mathrm{e}}\right)\mathbf{S}\right)^{\dagger} \\
&= \left(\mathbf{S}\left(\mathbf{Z}_{\mathrm{c}}\mathbf{K}_{\mathrm{c}}\mathbf{Z}_{\mathrm{c}}^{\mathrm{T}} + \mathbf{I}_n\delta_{\mathrm{e}}\right)\mathbf{S} + \sum_{l=1}^{L}\mathbf{S}\tilde{\mathbf{W}}_{\mathrm{r}_i,l}\tilde{\mathbf{\Gamma}}_{\mathrm{r}_i,l}\tilde{\mathbf{W}}_{\mathrm{r}_i,l}^{\mathrm{T}}\mathbf{S}\right)^{\dagger} \\
&= \left(\mathbf{U}_{\mathrm{L}}\left(\mathbf{\Lambda}_{\mathrm{c}} + \mathbf{I}_{n-p}\delta_{\mathrm{e}}\right)\mathbf{U}_{\mathrm{L}}^{\mathrm{T}} + \sum_{l=1}^{L}\mathbf{U}_{\mathrm{L}}\mathbf{U}_{\mathrm{L}}^{\mathrm{T}}\tilde{\mathbf{W}}_{\mathrm{r}_i,l}\tilde{\mathbf{\Gamma}}_{\mathrm{r}_i,l}\tilde{\mathbf{W}}_{\mathrm{r}_i,l}^{\mathrm{T}}\mathbf{U}_{\mathrm{L}}\mathbf{U}_{\mathrm{L}}^{\mathrm{T}}\right)^{\dagger} \\
&= \left(\mathbf{U}_{\mathrm{L}}\left(\mathbf{\Lambda}_{\mathrm{c}} + \mathbf{I}_{n-p}\delta_{\mathrm{e}} + \sum_{l=1}^{L}\mathbf{U}_{\mathrm{L}}^{\mathrm{T}}\tilde{\mathbf{W}}_{\mathrm{r}_i,l}\tilde{\mathbf{\Gamma}}_{\mathrm{r}_i,l}\tilde{\mathbf{W}}_{\mathrm{r}_i,l}^{\mathrm{T}}\mathbf{U}_{\mathrm{L}}\right)\mathbf{U}_{\mathrm{L}}^{\mathrm{T}}\right)^{\dagger} \\
&= \mathbf{U}_{\mathrm{L}}\left(\mathbf{\Lambda}_{\mathrm{c}} + \mathbf{I}_{n-p}\delta_{\mathrm{e}} + \sum_{l=1}^{L}\mathbf{U}_{\mathrm{L}}^{\mathrm{T}}\tilde{\mathbf{W}}_{\mathrm{r}_i,l}\tilde{\mathbf{\Gamma}}_{\mathrm{r}_i,l}\tilde{\mathbf{W}}_{\mathrm{r}_i,l}^{\mathrm{T}}\mathbf{U}_{\mathrm{L}}\right)^{-1}\mathbf{U}_{\mathrm{L}}^{\mathrm{T}} \\
&= \mathbf{U}_{\mathrm{L}}\left(\mathbf{\Lambda}_{\mathrm{c}} + \mathbf{I}_{n-p}\delta_{\mathrm{e}} + \sum_{l=1}^{L-1}\tilde{\mathbf{W}}_{\mathrm{r}_i,l}\tilde{\mathbf{\Gamma}}_{\mathrm{r}_i,l}\tilde{\mathbf{W}}_{\mathrm{r}_i,l}^{\mathrm{T}}\right)^{-1}\mathbf{U}_{\mathrm{L}}^{\mathrm{T}} \\
&\quad - \mathbf{U}_{\mathrm{L}}\left(\mathbf{\Lambda}_{\mathrm{c}} + \mathbf{I}_{n-p}\delta_{\mathrm{e}} + \sum_{l=1}^{L-1}\tilde{\mathbf{W}}_{\mathrm{r}_i,l}\tilde{\mathbf{\Gamma}}_{\mathrm{r}_i,l}\tilde{\mathbf{W}}_{\mathrm{r}_i,l}^{\mathrm{T}}\right)^{-1}\mathbf{U}_{\mathrm{L}}^{\mathrm{T}}\tilde{\mathbf{W}}_{\mathrm{r}_i,L} \\
&\quad \cdot \left(\tilde{\mathbf{\Gamma}}_{\mathrm{r}_i,L}^{-1} + \tilde{\mathbf{W}}_{\mathrm{r}_i,L}^{\mathrm{T}}\mathbf{U}_{\mathrm{L}}\left(\mathbf{\Lambda}_{\mathrm{c}} + \mathbf{I}_{n-p}\delta_{\mathrm{e}} + \sum_{l=1}^{L-1}\tilde{\mathbf{W}}_{\mathrm{r}_i,l}\tilde{\mathbf{\Gamma}}_{\mathrm{r}_i,l}\tilde{\mathbf{W}}_{\mathrm{r}_i,l}^{\mathrm{T}}\right)^{-1}\mathbf{U}_{\mathrm{L}}^{\mathrm{T}}\tilde{\mathbf{W}}_{\mathrm{r}_i,L}\right)^{-1} \\
&\quad \cdot \tilde{\mathbf{W}}_{\mathrm{r}_i,L}^{\mathrm{T}}\mathbf{U}_{\mathrm{L}}\left(\mathbf{\Lambda}_{\mathrm{c}} + \mathbf{I}_{n-p}\delta_{\mathrm{e}} + \sum_{l=1}^{L-1}\tilde{\mathbf{W}}_{\mathrm{r}_i,l}\tilde{\mathbf{\Gamma}}_{\mathrm{r}_i,l}\tilde{\mathbf{W}}_{\mathrm{r}_i,l}^{\mathrm{T}}\right)^{-1}\mathbf{U}_{\mathrm{L}}^{\mathrm{T}} \\
&= \mathbf{Q}^{(L-1)} - \mathbf{Q}^{(L-1)}\tilde{\mathbf{W}}_{\mathrm{r}_i,L}\left(\tilde{\mathbf{\Gamma}}_{\mathrm{r}_i,L}^{-1} + \tilde{\mathbf{W}}_{\mathrm{r}_i,L}^{\mathrm{T}}\mathbf{Q}^{(L-1)}\tilde{\mathbf{W}}_{\mathrm{r}_i,L}\right)^{-1}\tilde{\mathbf{W}}_{\mathrm{r}_i,L}^{\mathrm{T}}\mathbf{Q}^{(L-1)}
\end{aligned}
\tag{18}
$$

Here, we use Proposition 6, 5, 2, 7 and the Woodbury identity, and we define $\mathbf{Q}^{(L)}$ as the $\mathbf{Q}$ matrix for the $L$ random effects of the SNP-set, so $\mathbf{Q}^{(L)} = \mathbf{U}_{\mathrm{L}}\left(\mathbf{\Lambda}_{\mathrm{c}} + \mathbf{I}_{n-p}\delta_{\mathrm{e}} + \sum_{l=1}^{L}\mathbf{U}_{\mathrm{L}}^{\mathrm{T}}\tilde{\mathbf{W}}_{\mathrm{r}_i,l}\tilde{\mathbf{\Gamma}}_{\mathrm{r}_i,l}\tilde{\mathbf{W}}_{\mathrm{r}_i,l}^{\mathrm{T}}\mathbf{U}_{\mathrm{L}}\right)^{-1}\mathbf{U}_{\mathrm{L}}$. To compute $\mathbf{Q}^{(L)}$, we set $\mathbf{Q}^{(0)} = \mathbf{U}_{\mathrm{L}}\left(\mathbf{\Lambda}_{\mathrm{c}} + \mathbf{I}_{n-p}\delta_{\mathrm{e}}\right)$ first, and repeat Eq. (18) for $L$ steps. The computation of $\mathbf{Q}^{(L)}$ will be an $O(n^2\sum_{l=1}^{L}k_l)$ operation and it is much efficient if all $k_l \ll n$ where $k_l$ is the rank of $\tilde{\mathbf{W}}_{\mathrm{r}_i,l}$.

*Update of the determinant* $\log\left|\mathbf{Q}^{(L)}\right|_{+}$
We also extend Eq. (9) to

$$\begin{aligned}\log\left|\mathbf{Q}^{(L)}\right|_{+} &= -\log\left|\mathbf{U}_{\mathrm{L}}^{\mathrm{T}}\mathbf{H}\mathbf{U}_{\mathrm{L}}\right| \\ &= -\log\left|\mathbf{I}_{n-p}\delta_{\mathrm{e}} + \mathbf{\Lambda}_{\mathrm{c}} + \sum_{l=1}^{L}\mathbf{U}_{\mathrm{L}}^{\mathrm{T}}\tilde{\mathbf{W}}_{\mathrm{r}_i,l}\tilde{\mathbf{\Gamma}}_{\mathrm{r}_i,l}\tilde{\mathbf{W}}_{\mathrm{r}_i,l}^{\mathrm{T}}\mathbf{U}_{\mathrm{L}}\right| \\ &= -\log\left|\mathbf{I}_{n-p}\delta_{\mathrm{e}} + \mathbf{\Lambda}_{\mathrm{c}} + \sum_{l=1}^{L-1}\mathbf{A}^{(l)}\mathbf{B}^{(l)} + \mathbf{A}^{(L)}\mathbf{B}^{(L)}\right| \\ &= -\log\left|\mathbf{I}_{n-p}\delta_{\mathrm{e}} + \mathbf{\Lambda}_{\mathrm{c}} + \sum_{l=1}^{L-1}\mathbf{A}^{(l)}\mathbf{B}^{(l)}\right| \\ &\quad -\log\left|\mathbf{I}_{n-p} + \left(\mathbf{I}_{n-p}\delta_{\mathrm{e}} + \mathbf{\Lambda}_{\mathrm{c}} + \sum_{l=1}^{L-1}\mathbf{A}^{(l)}\mathbf{B}^{(l)}\right)^{-1}\mathbf{A}^{(L)}\mathbf{B}^{(L)}\right| \\ &= \log\left|\mathbf{Q}^{(L-1)}\right|_{+} - \log\left|\mathbf{I}_{k_L} + \mathbf{B}^{(L)}\mathbf{Q}^{(L-1)}\mathbf{A}^{(L)}\right| \end{aligned} \tag{19}$$

Here, we used Proposition 5, 6, 7 and the Sylvester's determinant identity, and we also define a $n-p \times k_L$ matrix $\mathbf{A}^{(L)} = \mathbf{U}_{\mathrm{L}}^{\mathrm{T}}\tilde{\mathbf{W}}_{\mathrm{r}_i,L}\tilde{\mathbf{\Gamma}}_{\mathrm{r}_i,L}$ and a $k \times n-p$ matrix $\mathbf{B}^{(L)} = \tilde{\mathbf{W}}_{\mathrm{r}_i,L}^{\mathrm{T}}\mathbf{U}_{\mathrm{L}}$. Computation of $\log\left|\mathbf{Q}^{(L)}\right|_{+}$ can be realized by repeating Eq. (19) after setting $\log\left|\mathbf{Q}^{(0)}\right|_{+} = -\log\left|\mathbf{I}_{n-p}\delta_{\mathrm{e}} + \mathbf{\Lambda}_{\mathrm{c}}\right|$. Here, when calculating the second term of Eq. (19), the notes mentioned for Eq. (9) still exists in this case.

#### The LR test for dominance and epistatic effects

To test the dominance or epistatic effects of each SNP-set, it should be assumed that the term of additive effects of each SNP-set is included both in the null model and the alternative model. Therefore we should assume the model such as Eq. (16) for the alternative model in this case.

If we define $\mathbf{K}_{\mathrm{r}_i,\mathrm{d}}$ is the dominance genetic matrix of $\mathbf{W}_{\mathrm{r}_i}$, $\tilde{\mathbf{W}}_{\mathrm{r}_i,\mathrm{d}}$ ($\tilde{\mathbf{W}}_{\mathrm{r}_i}$ for the dominance effects) corresponds to

$$\tilde{\mathbf{W}}_{\mathrm{r}_i,\mathrm{d}} = \frac{\mathbf{Z}_{\mathrm{r}_i}\left(\mathbf{1}_{m_{\mathrm{r}_i}\times M_i} - \|\mathbf{W}_{\mathrm{r}_i}\|\right)}{2\cdot\sum_{m=1}^{M_i} p_m\left(1-p_m\right)\left(1-p_m\left(1-p_m\right)\right)}, \tag{20}$$

and $\tilde{\mathbf{\Gamma}}_{\mathrm{r}_i,\mathrm{d}}$ is

$$\tilde{\mathbf{\Gamma}}_{\mathrm{r}_i,\mathrm{d}} = \mathbf{I}_{M_i}\gamma_{\mathrm{r}_i,\mathrm{d}}, \tag{21}$$

where $\gamma_{\mathrm{r}_i,\mathrm{d}}$ is the weight for the dominance effect of $i_{\mathrm{th}}$ SNP-set to be estimated [12]. Therefore, to test the significance of dominance effects, compare the restricted log likelihood of the model including both additive and dominance effects such as Eq. (16) ($l = 2$) with that of the model including only additive effects such as Eq. (1).

For the random effects of epistatic effects between two SNP-sets consisting of the same number of SNPs $M$, $\tilde{\mathbf{W}}_{\mathrm{r}_{ij},\mathrm{aa}}$, $\tilde{\mathbf{W}}_{\mathrm{r}_{ij},\mathrm{ad}}$, $\tilde{\mathbf{W}}_{\mathrm{r}_{ij},\mathrm{da}}$ and $\tilde{\mathbf{W}}_{\mathrm{r}_{ij},\mathrm{dd}}$($\tilde{\mathbf{W}}_{\mathrm{r}_{ij}}$ for additive

$\times$ additive, additive $\times$ dominance, dominance $\times$ additive and dominance $\times$ dominance epistatic effects between the $i_{\text{th}}$ and the $j_{\text{th}}$ SNP-sets of interest respectively) correspond to

$$\tilde{\mathbf{W}}_{\mathrm{r}_{ij},\mathrm{aa}} = \tilde{\mathbf{W}}_{\mathrm{r}_i,\mathrm{a}} \circ \tilde{\mathbf{W}}_{\mathrm{r}_j,\mathrm{a}}, \tag{22}$$

$$\tilde{\mathbf{W}}_{\mathrm{r}_{ij},\mathrm{ad}} = \tilde{\mathbf{W}}_{\mathrm{r}_i,\mathrm{a}} \circ \tilde{\mathbf{W}}_{\mathrm{r}_j,\mathrm{d}}, \tag{23}$$

$$\tilde{\mathbf{W}}_{\mathrm{r}_{ij},\mathrm{da}} = \tilde{\mathbf{W}}_{\mathrm{r}_i,\mathrm{d}} \circ \tilde{\mathbf{W}}_{\mathrm{r}_j,\mathrm{a}}, \tag{24}$$

$$\tilde{\mathbf{W}}_{\mathrm{r}_{ij},\mathrm{dd}} = \tilde{\mathbf{W}}_{\mathrm{r}_i,\mathrm{d}} \circ \tilde{\mathbf{W}}_{\mathrm{r}_j,\mathrm{d}}, \tag{25}$$

where $\tilde{\mathbf{W}}_{\mathrm{r}_i,\mathrm{a}}$ is defined by Eq. (10) and $\tilde{\mathbf{W}}_{\mathrm{r}_j,\mathrm{d}}$ is defined by Eq. (20) [13, 12]. On the other hand, correponding $\tilde{\mathbf{\Gamma}}_{\mathrm{r}_{ij},\mathrm{aa}}$, $\tilde{\mathbf{\Gamma}}_{\mathrm{r}_{ij},\mathrm{ad}}$, $\tilde{\mathbf{\Gamma}}_{\mathrm{r}_{ij},\mathrm{da}}$ and $\tilde{\mathbf{\Gamma}}_{\mathrm{r}_{ij},\mathrm{dd}}$ are

$$\tilde{\mathbf{\Gamma}}_{\mathrm{r}_{ij},\mathrm{aa}} = \mathbf{I}_M \gamma_{\mathrm{r}_{ij},\mathrm{aa}}, \tag{26}$$

$$\tilde{\mathbf{\Gamma}}_{\mathrm{r}_{ij},\mathrm{ad}} = \mathbf{I}_M \gamma_{\mathrm{r}_{ij},\mathrm{ad}}, \tag{27}$$

$$\tilde{\mathbf{\Gamma}}_{\mathrm{r}_{ij},\mathrm{da}} = \mathbf{I}_M \gamma_{\mathrm{r}_{ij},\mathrm{da}}, \tag{28}$$

$$\tilde{\mathbf{\Gamma}}_{\mathrm{r}_{ij},\mathrm{dd}} = \mathbf{I}_M \gamma_{\mathrm{r}_{ij},\mathrm{dd}}, \tag{29}$$

where $\gamma_{\mathrm{r}_{ij},\mathrm{aa}}$, $\gamma_{\mathrm{r}_{ij},\mathrm{ad}}$, $\gamma_{\mathrm{r}_{ij},\mathrm{da}}$ and $\gamma_{\mathrm{r}_{ij},\mathrm{dd}}$ are weights for each epistatic effects to be estimated. In this case, to test the significance of these epistatic effects, compare the restricted log likelihood of the model including additive and dominance effects of two SNP-set and four epistatic effects such as Eq. (16) ($l = 8$) with that of the model including only additive and dominance effects of two SNP-set such as Eq. (16) ($\mathrm{l} = 4$).

### Propositions

**Proposition 1** $\mathbf{SX} = \mathbf{0}$, $\mathbf{PX} = \mathbf{0}$ *and* $\mathbf{U}_{\mathrm{L}}^{\mathrm{T}}\mathbf{X} = \mathbf{0}$.

*Proof.*

$$\begin{aligned}
\mathbf{SX} &= \mathbf{X} - \mathbf{X}\left(\mathbf{X}^{\mathrm{T}}\mathbf{X}\right)^{-1}\mathbf{X}^{\mathrm{T}}\mathbf{X} = \mathbf{0},\\
\mathbf{PX} &= \mathbf{X} - \mathbf{X}\left(\mathbf{X}^{\mathrm{T}}\mathbf{H}^{-1}\mathbf{X}\right)^{-1}\mathbf{X}^{\mathrm{T}}\mathbf{H}^{-1}\mathbf{X} = \mathbf{0},
\end{aligned}$$

and because $\mathbf{PX} = \mathbf{0}$, $\mathbf{P}_{\mathrm{L}}\mathbf{X} = \mathbf{0}$. Therefore, $\mathbf{U}_{\mathrm{L}}^{\mathrm{T}}\mathbf{X} = \mathbf{0}$. □

**Proposition 2** $\mathbf{S}^2 = \mathbf{S}$.

*Proof.*

$$\begin{aligned}
\mathbf{S}^2 &= \left(\mathbf{I}_n - \mathbf{X}\left(\mathbf{X}^{\mathrm{T}}\mathbf{X}\right)^{-1}\mathbf{X}^{\mathrm{T}}\right)\left(\mathbf{I}_n - \mathbf{X}\left(\mathbf{X}^{\mathrm{T}}\mathbf{X}\right)^{-1}\mathbf{X}^{\mathrm{T}}\right),\\
&= \mathbf{I}_n - 2\cdot\mathbf{X}\left(\mathbf{X}^{\mathrm{T}}\mathbf{X}\right)^{-1}\mathbf{X}^{\mathrm{T}} + \mathbf{X}\left(\mathbf{X}^{\mathrm{T}}\mathbf{X}\right)^{-1}\mathbf{X}^{\mathrm{T}}\mathbf{X}\left(\mathbf{X}^{\mathrm{T}}\mathbf{X}\right)^{-1}\mathbf{X}^{\mathrm{T}}\\
&= \mathbf{I}_n - \mathbf{X}\left(\mathbf{X}^{\mathrm{T}}\mathbf{X}\right)^{-1}\mathbf{X}^{\mathrm{T}}\\
&= \mathbf{S}
\end{aligned}$$

This characteristic is called idempotent. □

**Proposition 3** $\mathbf{PS} = \mathbf{P}$ *and* $\mathbf{SP} = \mathbf{S}$.

*Proof.*

$$\begin{aligned}
\mathbf{PS} &= \left(\mathbf{I}_n - \mathbf{X}\left(\mathbf{X}^{\mathrm{T}}\mathbf{V}^{-1}\mathbf{X}\right)^{-1}\mathbf{X}^{\mathrm{T}}\mathbf{V}^{-1}\right)\left(\mathbf{I}_n - \mathbf{X}\left(\mathbf{X}^{\mathrm{T}}\mathbf{X}\right)^{-1}\mathbf{X}^{\mathrm{T}}\right)\\
&= \mathbf{I}_n - \mathbf{X}\left(\mathbf{X}^{\mathrm{T}}\mathbf{V}^{-1}\mathbf{X}\right)^{-1}\mathbf{X}^{\mathrm{T}}\mathbf{V}^{-1} - \mathbf{X}\left(\mathbf{X}^{\mathrm{T}}\mathbf{X}\right)^{-1}\mathbf{X}^{\mathrm{T}}\\
&\quad + \mathbf{X}\left(\mathbf{X}^{\mathrm{T}}\mathbf{V}^{-1}\mathbf{X}\right)^{-1}\mathbf{X}^{\mathrm{T}}\mathbf{V}^{-1}\mathbf{X}\left(\mathbf{X}^{\mathrm{T}}\mathbf{X}\right)^{-1}\mathbf{X}^{\mathrm{T}}\\
&= \mathbf{I}_n - \mathbf{X}\left(\mathbf{X}^{\mathrm{T}}\mathbf{V}^{-1}\mathbf{X}\right)^{-1}\mathbf{X}^{\mathrm{T}}\mathbf{V}^{-1}\\
&= \mathbf{P},
\end{aligned}$$

and

$$\begin{aligned}
\mathbf{SP} &= \left(\mathbf{I}_n - \mathbf{X}\left(\mathbf{X}^{\mathrm{T}}\mathbf{X}\right)^{-1}\mathbf{X}^{\mathrm{T}}\right)\left(\mathbf{I}_n - \mathbf{X}\left(\mathbf{X}^{\mathrm{T}}\mathbf{V}^{-1}\mathbf{X}\right)^{-1}\mathbf{X}^{\mathrm{T}}\mathbf{V}^{-1}\right)\\
&= \mathbf{I}_n - \mathbf{X}\left(\mathbf{X}^{\mathrm{T}}\mathbf{V}^{-1}\mathbf{X}\right)^{-1}\mathbf{X}^{\mathrm{T}}\mathbf{V}^{-1} - \mathbf{X}\left(\mathbf{X}^{\mathrm{T}}\mathbf{X}\right)^{-1}\mathbf{X}^{\mathrm{T}}\\
&\quad + \mathbf{X}\left(\mathbf{X}^{\mathrm{T}}\mathbf{X}\right)^{-1}\mathbf{X}^{\mathrm{T}}\mathbf{X}\left(\mathbf{X}^{\mathrm{T}}\mathbf{V}^{-1}\mathbf{X}\right)^{-1}\mathbf{X}^{\mathrm{T}}\mathbf{V}^{-1}\\
&= \mathbf{I}_n - \mathbf{X}\left(\mathbf{X}^{\mathrm{T}}\mathbf{X}\right)^{-1}\mathbf{X}^{\mathrm{T}}\\
&= \mathbf{S}
\end{aligned}$$

□

**Proposition 4** $\mathbf{P}^2 = \mathbf{P}$.

*Proof.*

$$
\begin{aligned}
\mathbf{P}^2 &= \left(\mathbf{I}_n - \mathbf{X}\left(\mathbf{X}^{\mathrm{T}}\mathbf{V}^{-1}\mathbf{X}\right)^{-1}\mathbf{X}^{\mathrm{T}}\mathbf{V}^{-1}\right)\left(\mathbf{I}_n - \mathbf{X}\left(\mathbf{X}^{\mathrm{T}}\mathbf{V}^{-1}\mathbf{X}\right)^{-1}\mathbf{X}^{\mathrm{T}}\mathbf{V}^{-1}\right) \\
&= \mathbf{I}_n - 2 \times \mathbf{X}\left(\mathbf{X}^{\mathrm{T}}\mathbf{V}^{-1}\mathbf{X}\right)^{-1}\mathbf{X}^{\mathrm{T}}\mathbf{V}^{-1} \\
&\quad + \mathbf{X}\left(\mathbf{X}^{\mathrm{T}}\mathbf{V}^{-1}\mathbf{X}\right)^{-1}\mathbf{X}^{\mathrm{T}}\mathbf{V}^{-1}\mathbf{X}\left(\mathbf{X}^{\mathrm{T}}\mathbf{V}^{-1}\mathbf{X}\right)^{-1}\mathbf{X}^{\mathrm{T}}\mathbf{V}^{-1} \\
&= \mathbf{I}_n - \mathbf{X}\left(\mathbf{X}^{\mathrm{T}}\mathbf{V}^{-1}\mathbf{X}\right)^{-1}\mathbf{X}^{\mathrm{T}}\mathbf{V}^{-1} \\
&= \mathbf{P}
\end{aligned}
$$

□

**Proposition 5** $\mathbf{P}^{\mathrm{T}}\mathbf{H}^{-1}\mathbf{P} = (\mathbf{SHS})^{\dagger}$.

*Proof.*
From Proposition 3 and 4,

$$
\begin{aligned}
(\mathbf{SHS})\left(\mathbf{P}^{\mathrm{T}}\mathbf{H}^{-1}\mathbf{P}\right)(\mathbf{SHS}) &= \mathbf{SHP}^{\mathrm{T}}\mathbf{H}^{-1}\mathbf{PHS} \\
&= \mathbf{SPPHS} \\
&= \mathbf{SHS},
\end{aligned}
$$

and

$$
\begin{aligned}
\left(\mathbf{P}^{\mathrm{T}}\mathbf{H}^{-1}\mathbf{P}\right)(\mathbf{SHS})\left(\mathbf{P}^{\mathrm{T}}\mathbf{H}^{-1}\mathbf{P}\right) &= \mathbf{P}^{\mathrm{T}}\mathbf{H}^{-1}\mathbf{PHP}^{\mathrm{T}}\mathbf{H}^{-1}\mathbf{P} \\
&= \mathbf{P}^{\mathrm{T}}\mathbf{H}^{-1}\mathbf{PPP} \\
&= \mathbf{P}^{\mathrm{T}}\mathbf{H}^{-1}\mathbf{P}.
\end{aligned}
$$

Moreover, the product of $\mathbf{P}^{\mathrm{T}}\mathbf{H}^{-1}\mathbf{P}$ and $\mathbf{SHS}$ is is commutative and becomes a Hermitian (in this case, real symmetric) matrix.

$$
(\mathbf{SHS})\left(\mathbf{P}^{\mathrm{T}}\mathbf{H}^{-1}\mathbf{P}\right) = \left(\mathbf{P}^{\mathrm{T}}\mathbf{H}^{-1}\mathbf{P}\right)(\mathbf{SHS}) = \mathbf{S}
$$

Therefore, from the definition of Moore-Penrose peudo-inverse matrix,

$$
\mathbf{P}^{\mathrm{T}}\mathbf{H}^{-1}\mathbf{P} = (\mathbf{SHS})^{\dagger} = \mathbf{U}_{\mathrm{L}}\mathbf{\Lambda}_{L}^{-1}\mathbf{U}_{\mathrm{L}}^{\mathrm{T}}
$$

□

**Proposition 6** $\mathbf{Q} = \mathbf{P}^{\mathrm{T}}\mathbf{H}^{-1}\mathbf{P}$.

*Proof.*
By using Proposition 4,

$$
\mathbf{P}^{\mathrm{T}}\mathbf{H}^{-1}\mathbf{P} = \mathbf{H}^{-1}\mathbf{PP} = \mathbf{H}^{-1}\mathbf{P} = \mathbf{Q}
$$

□

**Proposition 7** $\mathbf{U}_{\mathrm{L}}\mathbf{U}_{\mathrm{L}}^{\mathrm{T}} = \mathbf{S}$.

*Proof.*
By using Proposition 5,

$$\begin{aligned}\mathbf{S} &= (\mathbf{SHS})\left(\mathbf{P}^{\mathrm{T}}\mathbf{H}^{-1}\mathbf{P}\right) = (\mathbf{SHS})\,(\mathbf{SHS})^{\dagger} \\ &= \mathbf{U}_{\mathrm{L}}\mathbf{\Lambda}_{L}\mathbf{U}_{\mathrm{L}}^{\mathrm{T}}\mathbf{U}_{\mathrm{L}}\mathbf{\Lambda}_{L}^{-1}\mathbf{U}_{\mathrm{L}}^{\mathrm{T}} \\ &= \mathbf{U}_{\mathrm{L}}\mathbf{U}_{\mathrm{L}}^{\mathrm{T}}\end{aligned}$$

□

**Proposition 8** $\log\left|\mathbf{P}_{\mathrm{L}}\mathbf{V}\mathbf{P}_{\mathrm{L}}^{\mathrm{T}}\right| = -\log\left|\mathbf{P}^{\mathrm{T}}\mathbf{V}^{-1}\mathbf{P}\right|_{+}$.

*Proof.*
First,

$$\mathbf{P}_{\mathrm{L}}\mathbf{V}\mathbf{P}_{\mathrm{L}}^{\mathrm{T}} = \mathbf{U}_{\mathrm{L}}^{\mathrm{T}}\mathbf{H}\mathbf{U}_{\mathrm{L}}\sigma_{\mathrm{c}}^{2} = \mathbf{\Lambda}_{L}\sigma_{\mathrm{c}}^{2},$$

because

$$\mathbf{U}_{\mathrm{L}}\mathbf{U}_{\mathrm{L}}^{\mathrm{T}}\mathbf{H}\mathbf{U}_{\mathrm{L}}\mathbf{U}_{\mathrm{L}}^{\mathrm{T}} = \mathbf{SHS} = \mathbf{U}_{\mathrm{L}}\mathbf{\Lambda}_{L}\mathbf{U}_{\mathrm{L}}^{\mathrm{T}},$$

from Proposition 7. Then, by using Proposition 5

$$\begin{aligned}\log\left|\mathbf{\Lambda}_{L}\right| &= -\log\left|\mathbf{\Lambda}_{L}^{-1}\right| \\ &= -\log\left|\mathbf{U}_{\mathrm{L}}\mathbf{\Lambda}_{L}^{-1}\mathbf{U}_{\mathrm{L}}^{\mathrm{T}}\right|_{+} \\ &= -\log\left|\mathbf{P}^{\mathrm{T}}\mathbf{H}^{-1}\mathbf{P}\right|_{+}\end{aligned}$$

Therefore,

$$\log\left|\mathbf{P}_{\mathrm{L}}\mathbf{V}\mathbf{P}_{\mathrm{L}}^{\mathrm{T}}\right| = \log\left|\mathbf{\Lambda}_{L}\sigma_{\mathrm{c}}^{2}\right| = -\log\left|\mathbf{P}^{\mathrm{T}}\mathbf{V}^{-1}\mathbf{P}\right|_{+}$$

□

**Proposition 9** $\mathbf{P}_{\mathrm{L}}^{\mathrm{T}}\left(\mathbf{P}_{\mathrm{L}}\mathbf{V}\mathbf{P}_{\mathrm{L}}^{\mathrm{T}}\right)^{-1}\mathbf{P}_{\mathrm{L}} = \mathbf{V}^{-1}\mathbf{P}$.

*Proof.* By using Proposition 8 and 6,

$$\begin{aligned}\mathbf{P}_{\mathrm{L}}^{\mathrm{T}}\left(\mathbf{P}_{\mathrm{L}}\mathbf{V}\mathbf{P}_{\mathrm{L}}^{\mathrm{T}}\right)^{-1}\mathbf{P}_{\mathrm{L}} &= \mathbf{U}_{\mathrm{L}}\left(\mathbf{U}_{\mathrm{L}}^{\mathrm{T}}\mathbf{V}\mathbf{U}_{\mathrm{L}}\right)^{-1}\mathbf{U}_{\mathrm{L}}^{\mathrm{T}} \\ &= \mathbf{U}_{\mathrm{L}}\left(\mathbf{\Lambda}_{L}\sigma_{\mathrm{c}}^{2}\right)^{-1}\mathbf{U}_{\mathrm{L}}^{\mathrm{T}} \\ &= \mathbf{P}^{\mathrm{T}}\mathbf{H}^{-1}\mathbf{P}/\sigma_{\mathrm{c}}^{2} \\ &= \mathbf{Q}/\sigma_{\mathrm{c}}^{2} = \mathbf{V}^{-1}\mathbf{P}\end{aligned}$$

□

**Proposition 10** $\frac{1}{2}\log|\mathbf{Q}|_{+} = -\frac{1}{2}\log|\mathbf{H}| + \frac{1}{2}\log\left|\mathbf{X}^{\mathrm{T}}\mathbf{X}\right| - \frac{1}{2}\log\left|\mathbf{X}^{\mathrm{T}}\mathbf{H}^{-1}\mathbf{X}\right|$.

*Proof.* By using Proposition 6, 5 and 7,

$$\begin{aligned}
|\mathbf{Q}|_{+} &= \left|\mathbf{P}^{\mathrm{T}}\mathbf{H}^{-1}\mathbf{P}\right|_{+} \\
&= |\mathbf{SHS}|_{+}^{-1} \\
&= |\mathbf{\Lambda}_{L}|^{-1} \cdot \left|\mathbf{X}^{\mathrm{T}}\mathbf{H}^{-1}\mathbf{X}\right| \cdot \left|\mathbf{X}^{\mathrm{T}}\mathbf{H}^{-1}\mathbf{X}\right|^{-1} \\
&= \left|\mathbf{U}_{\mathrm{L}}^{\mathrm{T}}\mathbf{H}\mathbf{U}_{\mathrm{L}}\right|^{-1} \cdot \left|\left(\mathbf{X}^{\mathrm{T}}\mathbf{H}^{-1}\mathbf{X}\right)^{-1}\mathbf{X}^{\mathrm{T}}\mathbf{H}^{-1}\mathbf{X}\left(\mathbf{X}^{\mathrm{T}}\mathbf{H}^{-1}\mathbf{X}\right)^{-1}\right|^{-1} \cdot \left|\mathbf{X}^{\mathrm{T}}\mathbf{H}^{-1}\mathbf{X}\right|^{-1} \\
&= \left|\mathbf{U}_{\mathrm{L}}^{\mathrm{T}}\mathbf{H}\mathbf{U}_{\mathrm{L}}\right|^{-1} \cdot \left|\left(\mathbf{X}^{\mathrm{T}}\mathbf{H}^{-1}\mathbf{X}\right)^{-1}\mathbf{X}^{\mathrm{T}}\mathbf{H}^{-1}\mathbf{H}\mathbf{H}^{-1}\mathbf{X}\left(\mathbf{X}^{\mathrm{T}}\mathbf{H}^{-1}\mathbf{X}\right)^{-1}\right|^{-1} \cdot \left|\mathbf{X}^{\mathrm{T}}\mathbf{H}^{-1}\mathbf{X}\right|^{-1} \\
&= \left|\begin{bmatrix}\mathbf{U}_{\mathrm{L}}^{\mathrm{T}}\mathbf{H}\mathbf{U}_{\mathrm{L}} & \mathbf{0} \\ \mathbf{0} & \left(\mathbf{X}^{\mathrm{T}}\mathbf{H}^{-1}\mathbf{X}\right)^{-1}\mathbf{X}^{\mathrm{T}}\mathbf{H}^{-1}\mathbf{H}\mathbf{H}^{-1}\mathbf{X}\left(\mathbf{X}^{\mathrm{T}}\mathbf{H}^{-1}\mathbf{X}\right)^{-1}\end{bmatrix}\right|^{-1} \cdot \left|\mathbf{X}^{\mathrm{T}}\mathbf{H}^{-1}\mathbf{X}\right|^{-1}
\end{aligned}$$

Here, we define $\mathbf{M} = \left(\mathbf{X}^{\mathrm{T}}\mathbf{H}^{-1}\mathbf{X}\right)^{-1}\mathbf{X}^{\mathrm{T}}\mathbf{H}^{-1}$ to shorten the notation.

$$= \left|\begin{bmatrix}\mathbf{U}_{\mathrm{L}}^{\mathrm{T}}\mathbf{H}\mathbf{U}_{\mathrm{L}} & \mathbf{U}_{\mathrm{L}}^{\mathrm{T}}\mathbf{H}\mathbf{M}^{\mathrm{T}} \\ \mathbf{M}\mathbf{H}\mathbf{U}_{\mathrm{L}} & \mathbf{M}\mathbf{H}\mathbf{M}^{\mathrm{T}}\end{bmatrix}\right|^{-1} \cdot \left|\mathbf{X}^{\mathrm{T}}\mathbf{H}^{-1}\mathbf{X}\right|^{-1}$$

$\mathbf{M}\mathbf{H}\mathbf{U}_{\mathrm{L}} = \mathbf{0}$ because $\mathbf{X}^{\mathrm{T}}\mathbf{U}_{\mathrm{L}} = \mathbf{0}$ (Proposition 1).

$$\begin{aligned}
&= \left|\begin{bmatrix}\mathbf{U}_{\mathrm{L}}^{\mathrm{T}} \\ \mathbf{M}\end{bmatrix}\mathbf{H}\begin{bmatrix}\mathbf{U}_{\mathrm{L}}^{\mathrm{T}} \\ \mathbf{M}\end{bmatrix}^{\mathrm{T}}\right|^{-1} \cdot \left|\mathbf{X}^{\mathrm{T}}\mathbf{H}^{-1}\mathbf{X}\right|^{-1} \\
&= |\mathbf{H}|^{-1} \cdot \left|\begin{bmatrix}\mathbf{U}_{\mathrm{L}}^{\mathrm{T}} \\ \mathbf{M}\end{bmatrix}\begin{bmatrix}\mathbf{U}_{\mathrm{L}}^{\mathrm{T}} \\ \mathbf{M}\end{bmatrix}^{\mathrm{T}}\right|^{-1} \cdot \left|\mathbf{X}^{\mathrm{T}}\mathbf{H}^{-1}\mathbf{X}\right|^{-1} \\
&= |\mathbf{H}|^{-1} \cdot \left|\begin{bmatrix}\mathbf{U}_{\mathrm{L}}^{\mathrm{T}}\mathbf{U}_{\mathrm{L}} & \mathbf{U}_{\mathrm{L}}^{\mathrm{T}}\mathbf{M}^{\mathrm{T}} \\ \mathbf{M}\mathbf{U}_{\mathrm{L}} & \mathbf{M}\mathbf{M}^{\mathrm{T}}\end{bmatrix}\right|^{-1} \cdot \left|\mathbf{X}^{\mathrm{T}}\mathbf{H}^{-1}\mathbf{X}\right|^{-1}
\end{aligned}$$

Using the well-known formula for the determinant of block-matrix,

$$\begin{aligned}
&= |\mathbf{H}|^{-1} \cdot \left|\mathbf{U}_{\mathrm{L}}^{\mathrm{T}}\mathbf{U}_{\mathrm{L}}\right|^{-1} \cdot \left|\mathbf{M}\mathbf{M}^{\mathrm{T}} - \mathbf{M}\mathbf{U}_{\mathrm{L}}\left(\mathbf{U}_{\mathrm{L}}^{\mathrm{T}}\mathbf{U}_{\mathrm{L}}\right)^{-1}\mathbf{U}_{\mathrm{L}}^{\mathrm{T}}\mathbf{M}^{\mathrm{T}}\right|^{-1} \cdot \left|\mathbf{X}^{\mathrm{T}}\mathbf{H}^{-1}\mathbf{X}\right|^{-1} \\
&= |\mathbf{H}|^{-1} \cdot \left|\mathbf{M}\mathbf{M}^{\mathrm{T}} - \mathbf{M}\mathbf{S}\mathbf{M}^{\mathrm{T}}\right|^{-1} \cdot \left|\mathbf{X}^{\mathrm{T}}\mathbf{H}^{-1}\mathbf{X}\right|^{-1} \\
&= |\mathbf{H}|^{-1} \cdot \left|\mathbf{M}\mathbf{X}\left(\mathbf{X}^{\mathrm{T}}\mathbf{X}\right)^{-1}\mathbf{X}^{\mathrm{T}}\mathbf{M}^{\mathrm{T}}\right|^{-1} \cdot \left|\mathbf{X}^{\mathrm{T}}\mathbf{H}^{-1}\mathbf{X}\right|^{-1} \\
&= |\mathbf{H}|^{-1} \cdot \left|\mathbf{X}^{\mathrm{T}}\mathbf{X}\right| \cdot \left|\mathbf{X}^{\mathrm{T}}\mathbf{H}^{-1}\mathbf{X}\right|^{-1}
\end{aligned}$$

Therefore,

$$\frac{1}{2}\log|\mathbf{Q}|_{+} = -\frac{1}{2}\log|\mathbf{H}| + \frac{1}{2}\log\left|\mathbf{X}^{\mathrm{T}}\mathbf{X}\right| - \frac{1}{2}\log\left|\mathbf{X}^{\mathrm{T}}\mathbf{H}^{-1}\mathbf{X}\right|$$

If we plus the both sides of the equation by $-(n-p)\log(\sigma_{\mathrm{c}}^{2})/2$, we get

$$\frac{1}{2}\log\left|\mathbf{Q}/\sigma_{\mathrm{c}}^{2}\right|_{+} = -\frac{1}{2}\log|\mathbf{V}| + \frac{1}{2}\log\left|\mathbf{X}^{\mathrm{T}}\mathbf{X}\right| - \frac{1}{2}\log\left|\mathbf{X}^{\mathrm{T}}\mathbf{V}^{-1}\mathbf{X}\right|,$$

which is also the frequently used form of the formula. □

**Author details**
