## Supplementary figures and images for "Haplotype-based genome wide association study using a novel SNP-set method : RAINBOW"

### Supplementary Figure 1

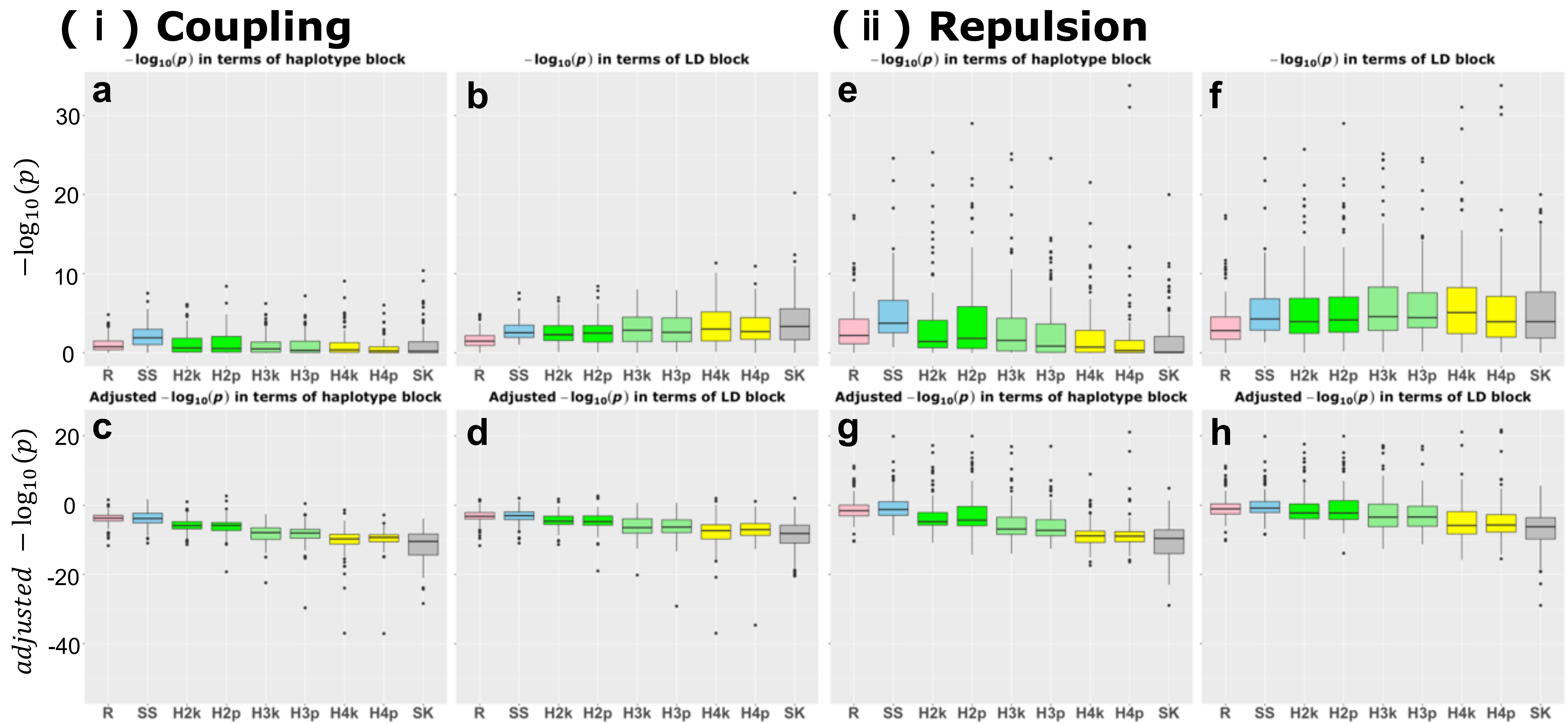

### Supplementary Figure 2

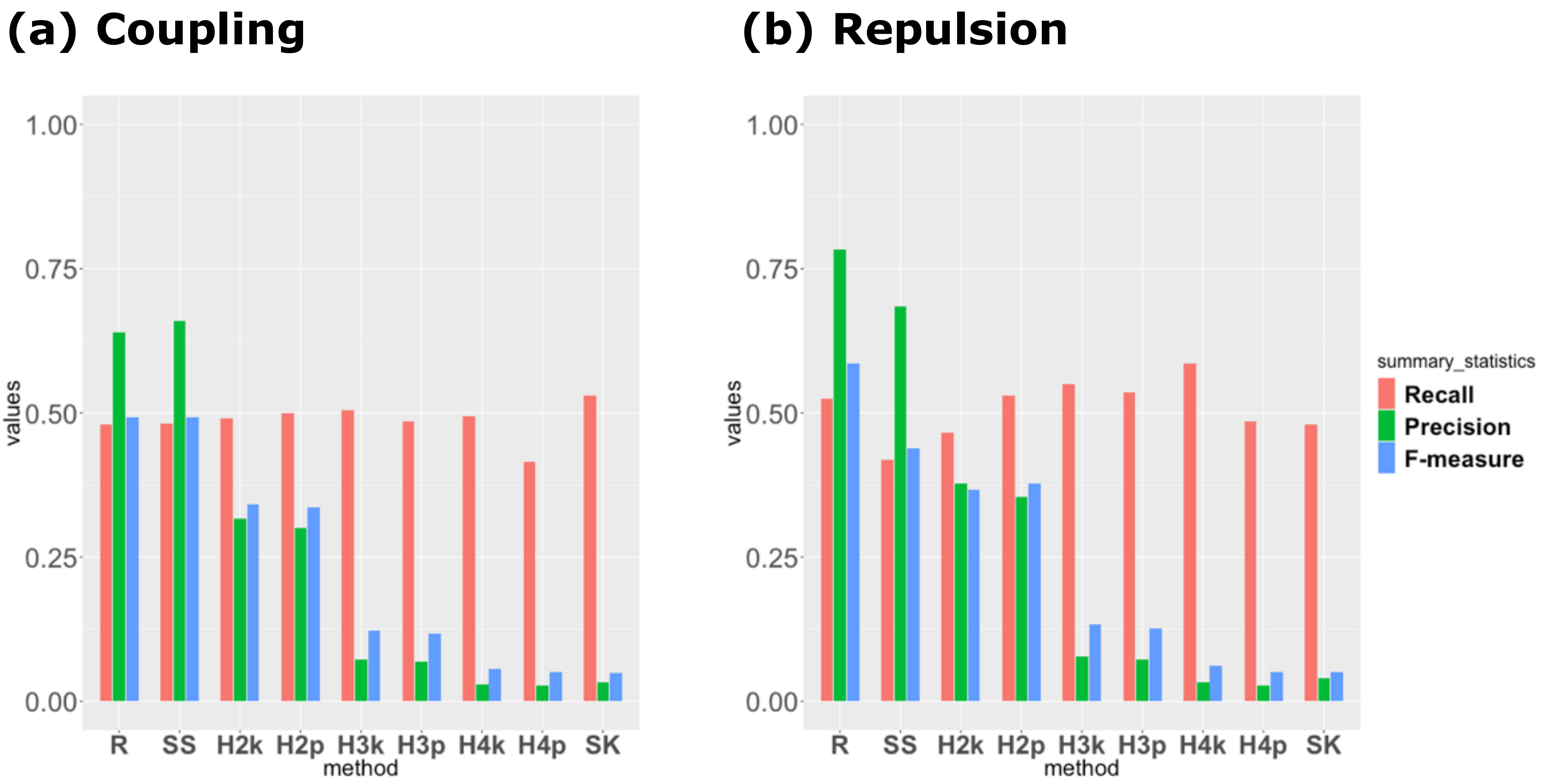

### Supplementary Figure 3

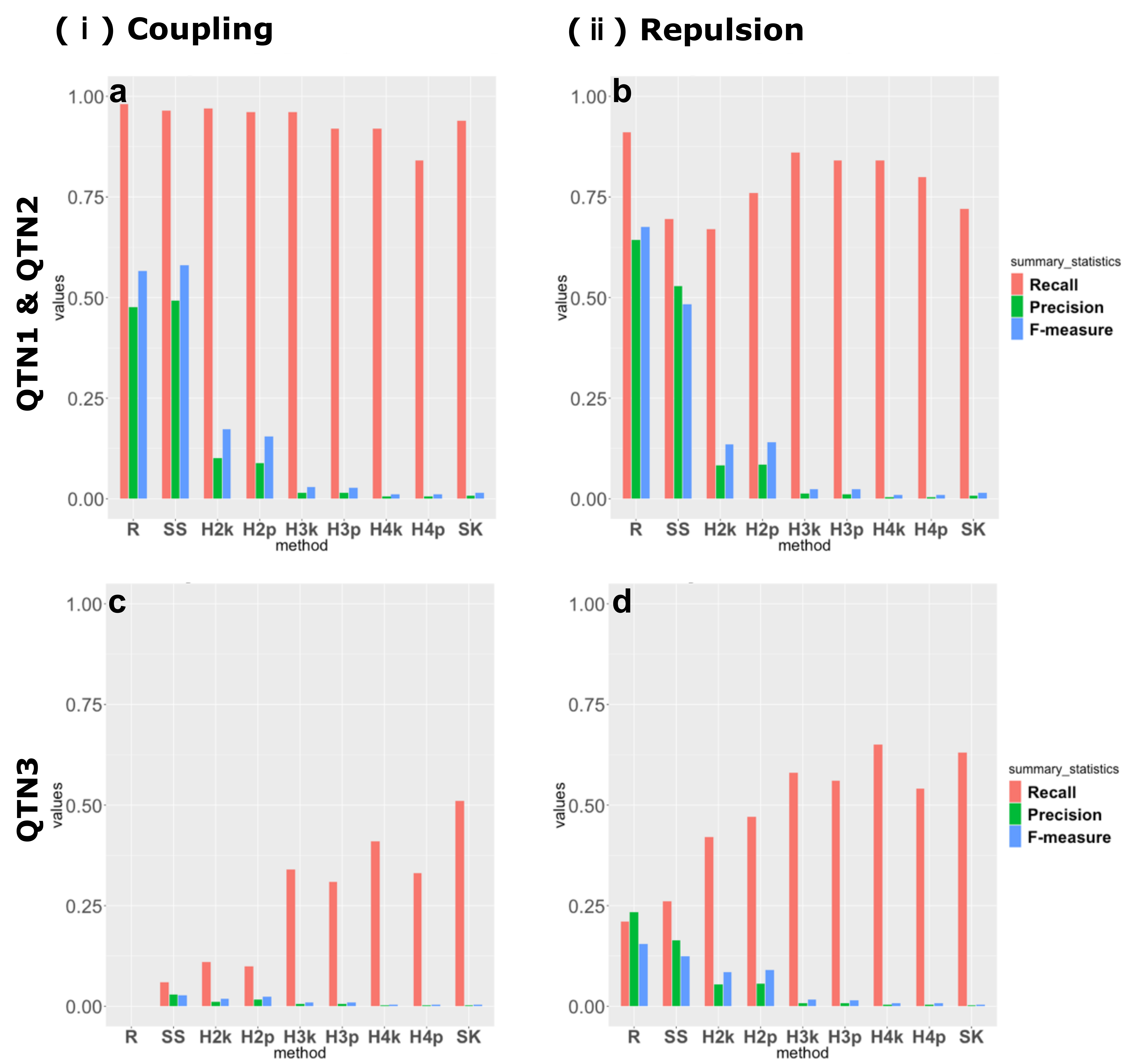

### Supplementary Figure 4

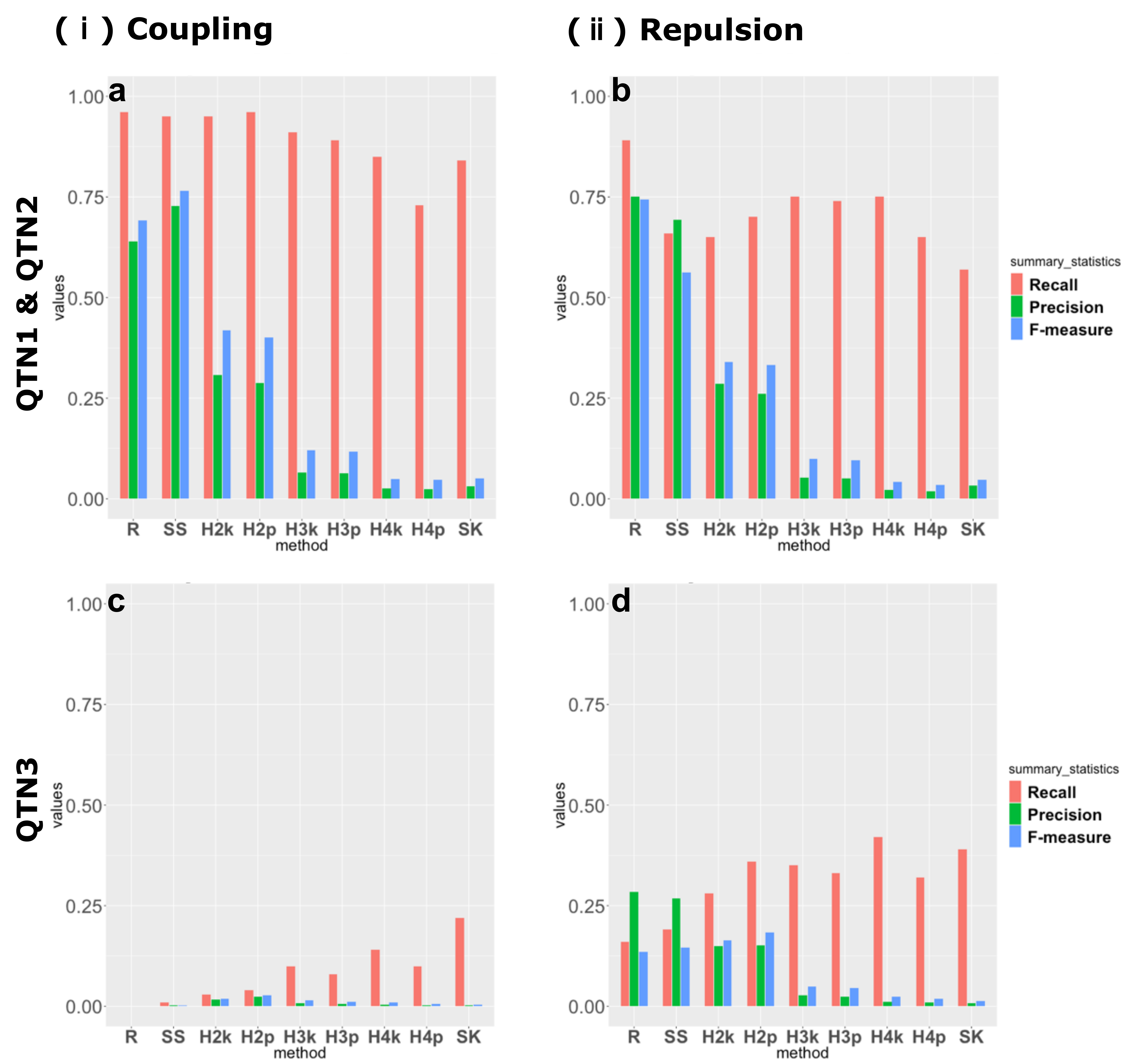

### Supplementary Figure 5

#### Iteration 40

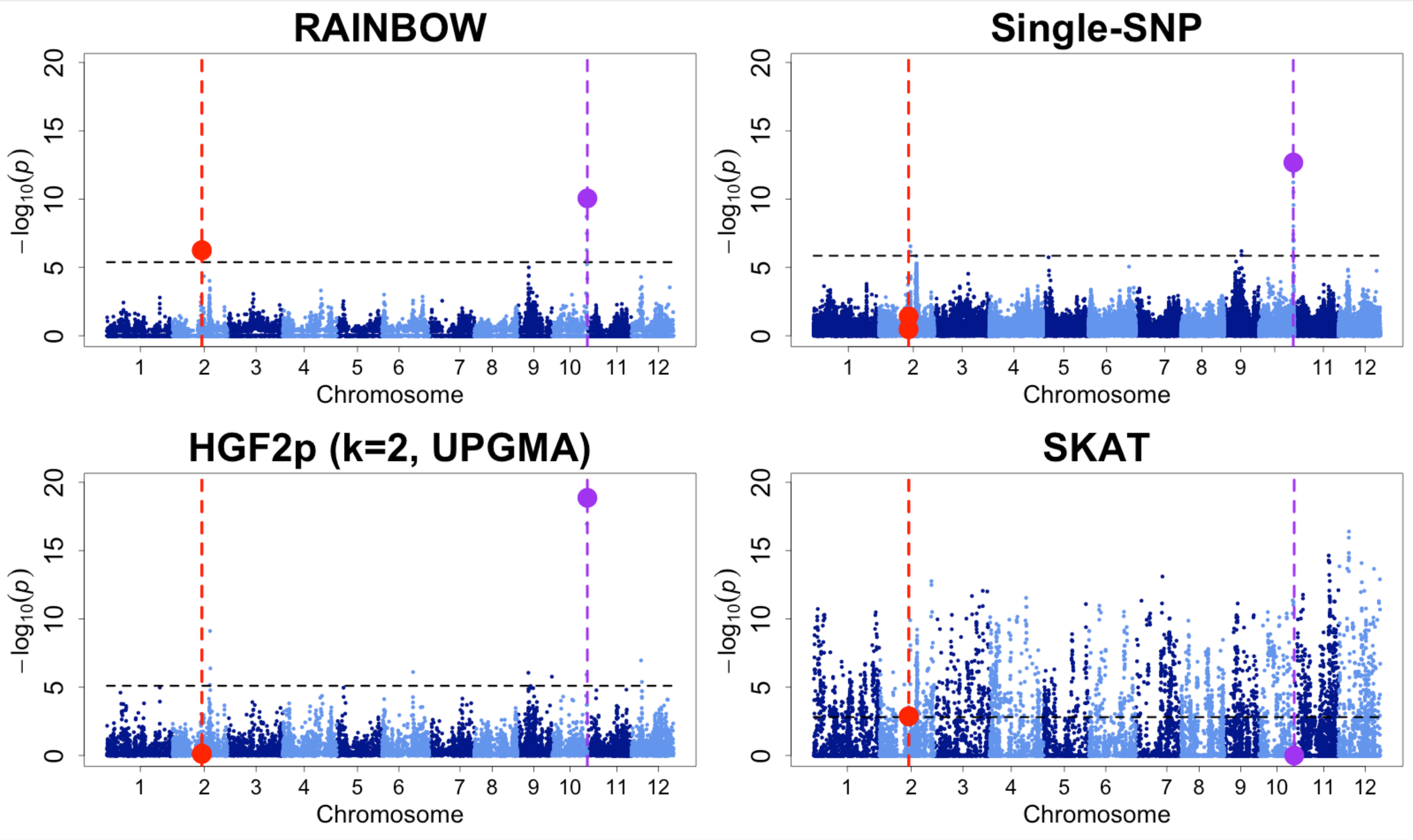

#### Iteration 43

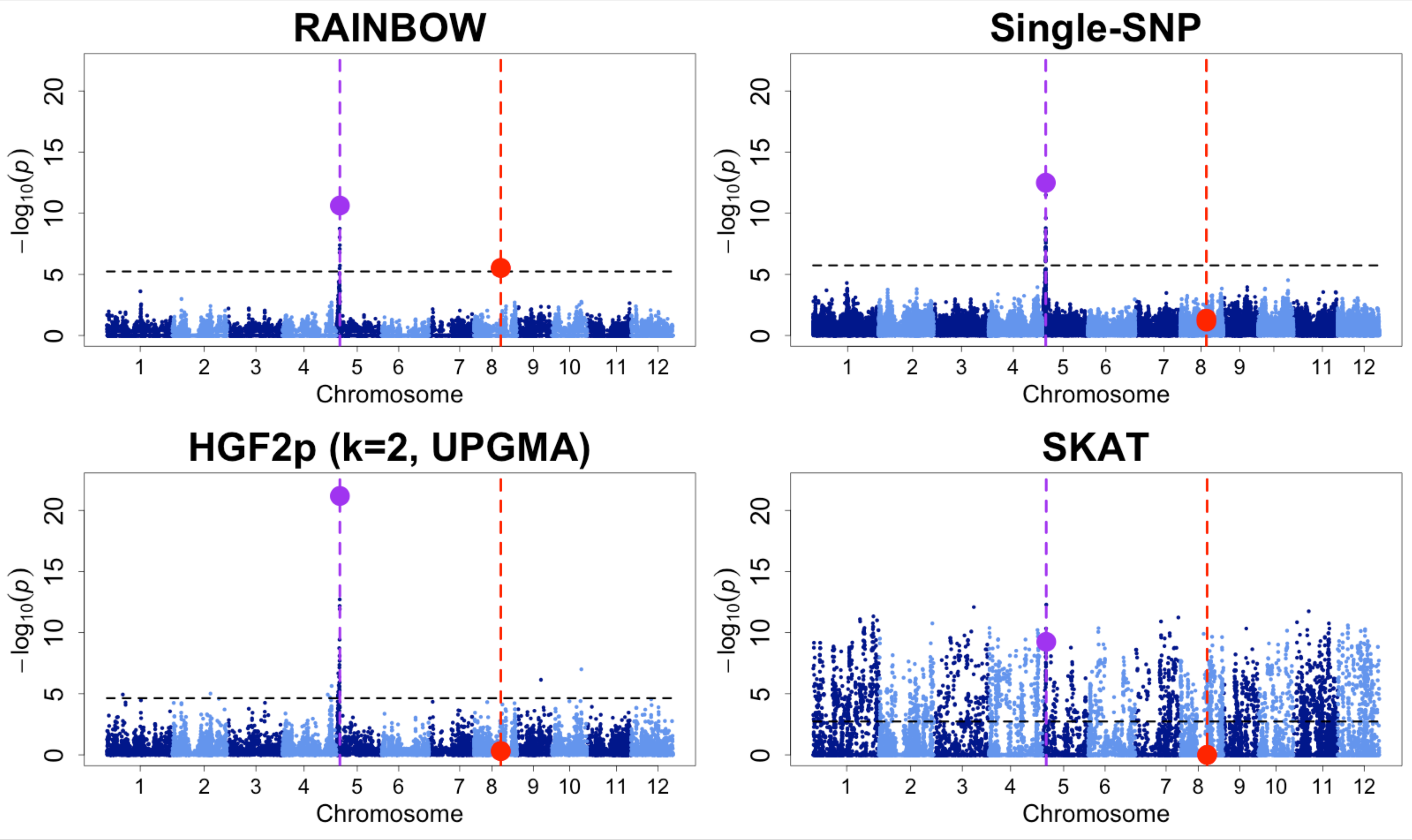

#### Iteration 64

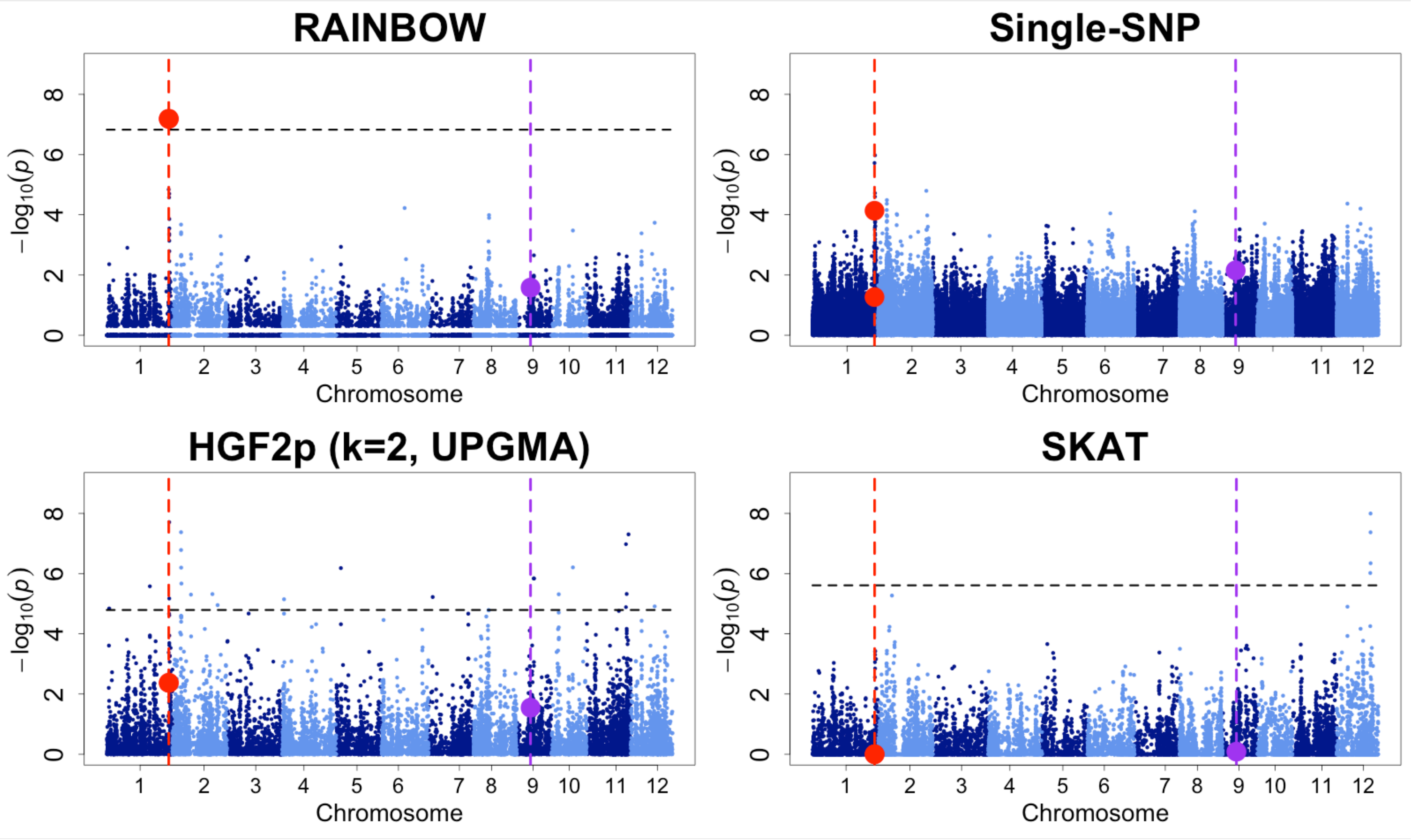

#### Iteration 69

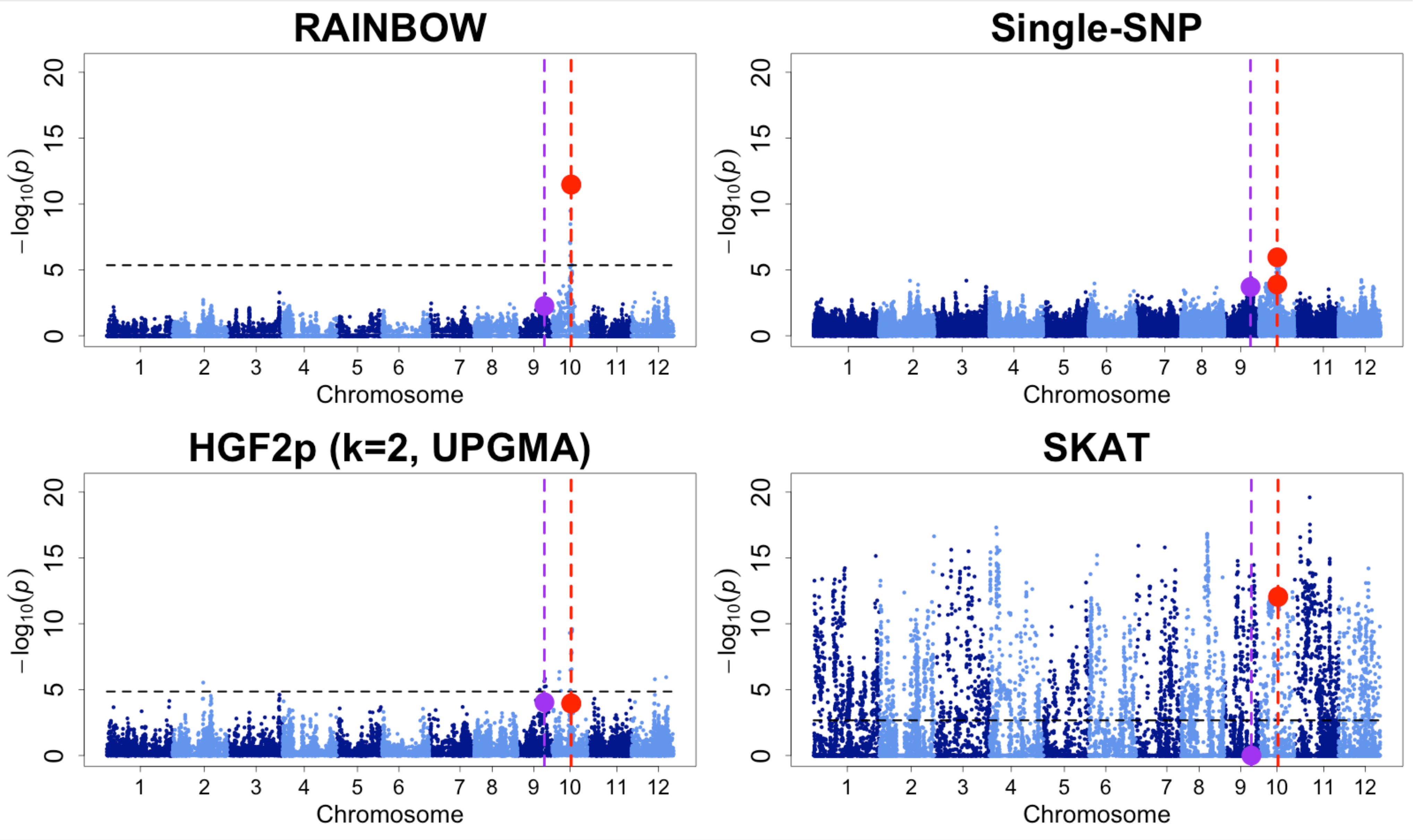

#### Iteration 81

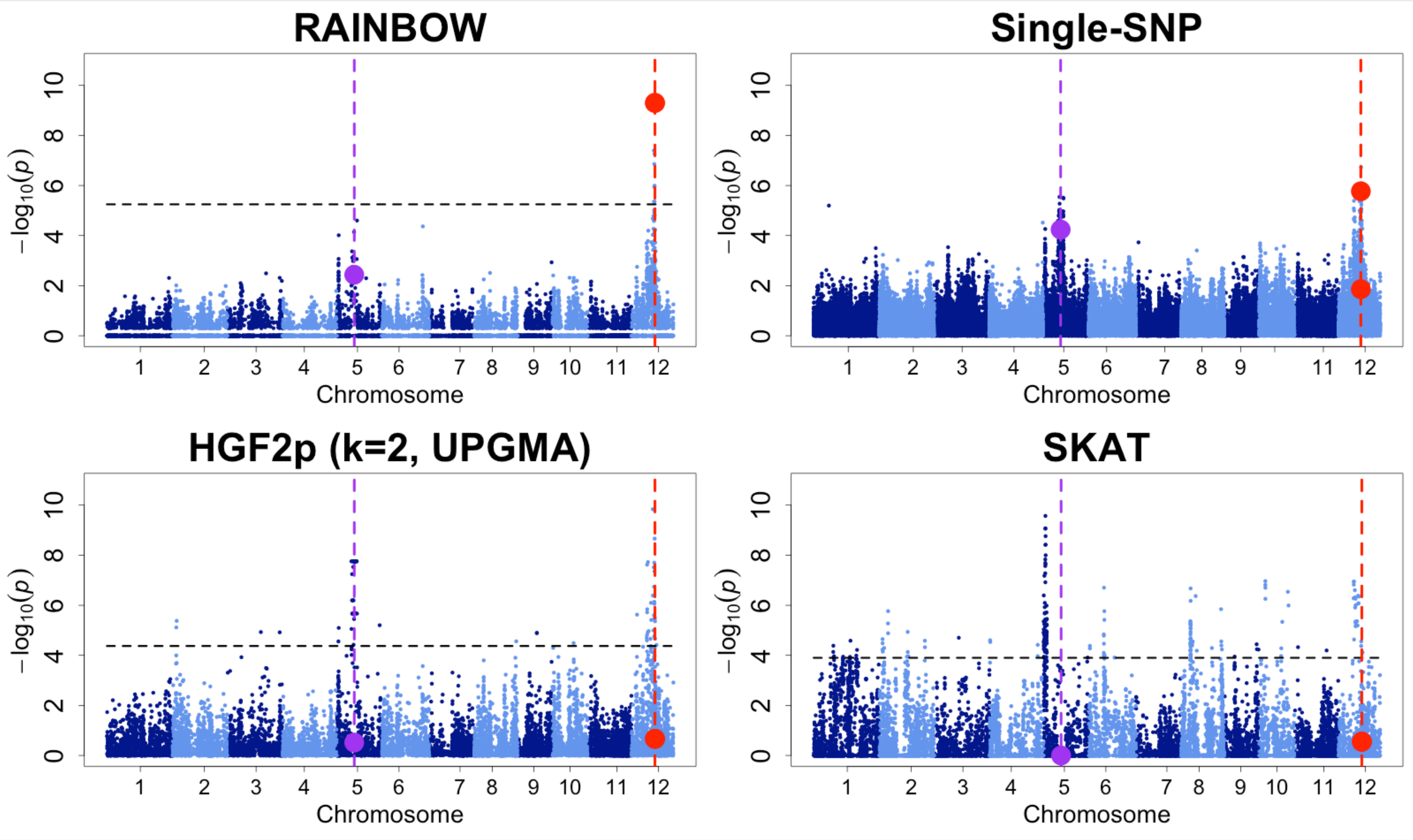

#### Iteration 85

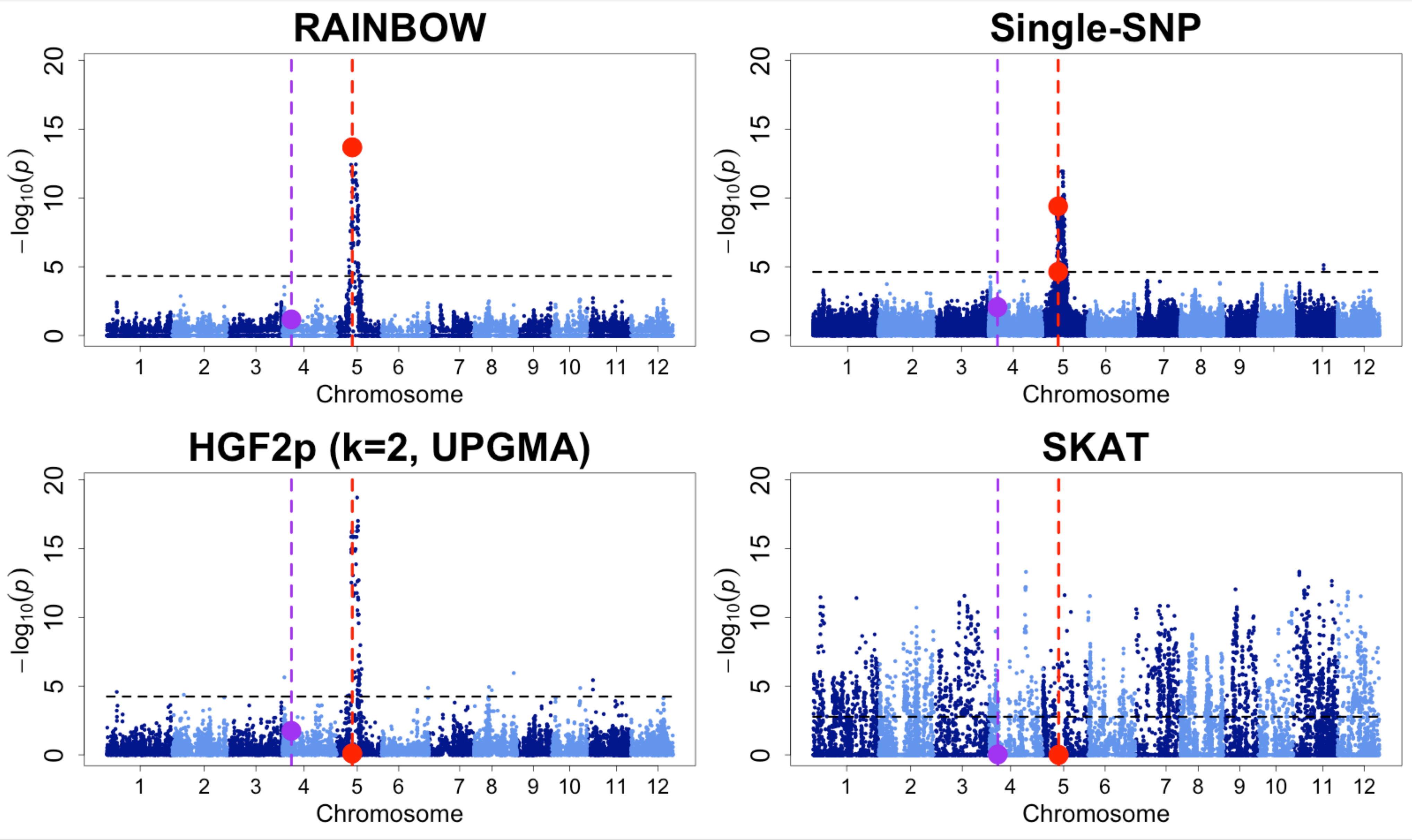
